# Optimal policy for multi-alternative decisions

**DOI:** 10.1101/595843

**Authors:** Satohiro Tajima, Jan Drugowitsch, Nisheet Patel, Alexandre Pouget

## Abstract

Every-day decisions frequently require choosing among multiple alternatives. Yet, the optimal policy for such decisions is unknown. Here we derive the normative policy for general multi-alternative decisions. This strategy requires evidence accumulation to nonlinear, time-dependent bounds, that trigger choices. A geometric symmetry in those boundaries allows the optimal strategy to be implemented by a simple neural circuit involving a normalization with fixed decision bounds and an urgency signal. The model captures several key features of the response of decision-making neurons as well as the increase in reaction time as a function of the number of alternatives, known as Hick’s law. In addition, we show that, in the presence of divisive normalization and internal variability, our model can account for several so called ‘irrational’ behaviors such as the similarity effect as well as the violation of both the independent irrelevant alternative principle and the regularity principle.

## Introduction

In a natural environment, choosing the best of multiple options is frequently critical for an organism’s survival. Such decisions are often value-based, in which case the reward is determined by the chosen item (such as when subjects chose between food items; Figure 1a), or perceptual, in which case subjects receive a fixed reward if they pick the correct option (Figure 1b). Compared to binary choice paradigms^1–3^, much less is known about the computational principles underlying decisions with more than two options^4^. Some studies have suggested that decisions among 3 or 4 options could be solved with coupled drift diffusion models^4–6^, which are optimal for binary choices^7^, but, as we’ll show, become sub-optimal once the number of choices grows beyond two. Another option for modelling such choices is to use “race models” (RMs). In RMs, the momentary choice preference is encoded by competing evidence accumulators, one per options, which trigger a choice as soon as one of them reaches a decision threshold (Figure 1c). Such standard RMs imply that both the races and the static decision criteria are independent across individual options. However, in contrast to race models, the nervous system features dynamic neural interactions across races, such as activity normalization^8,9^ and a global urgency signal^10^. Whether such coupled races are compatible with optimal decision policies for 3 or more choices is unknown.

**Figure 1.**
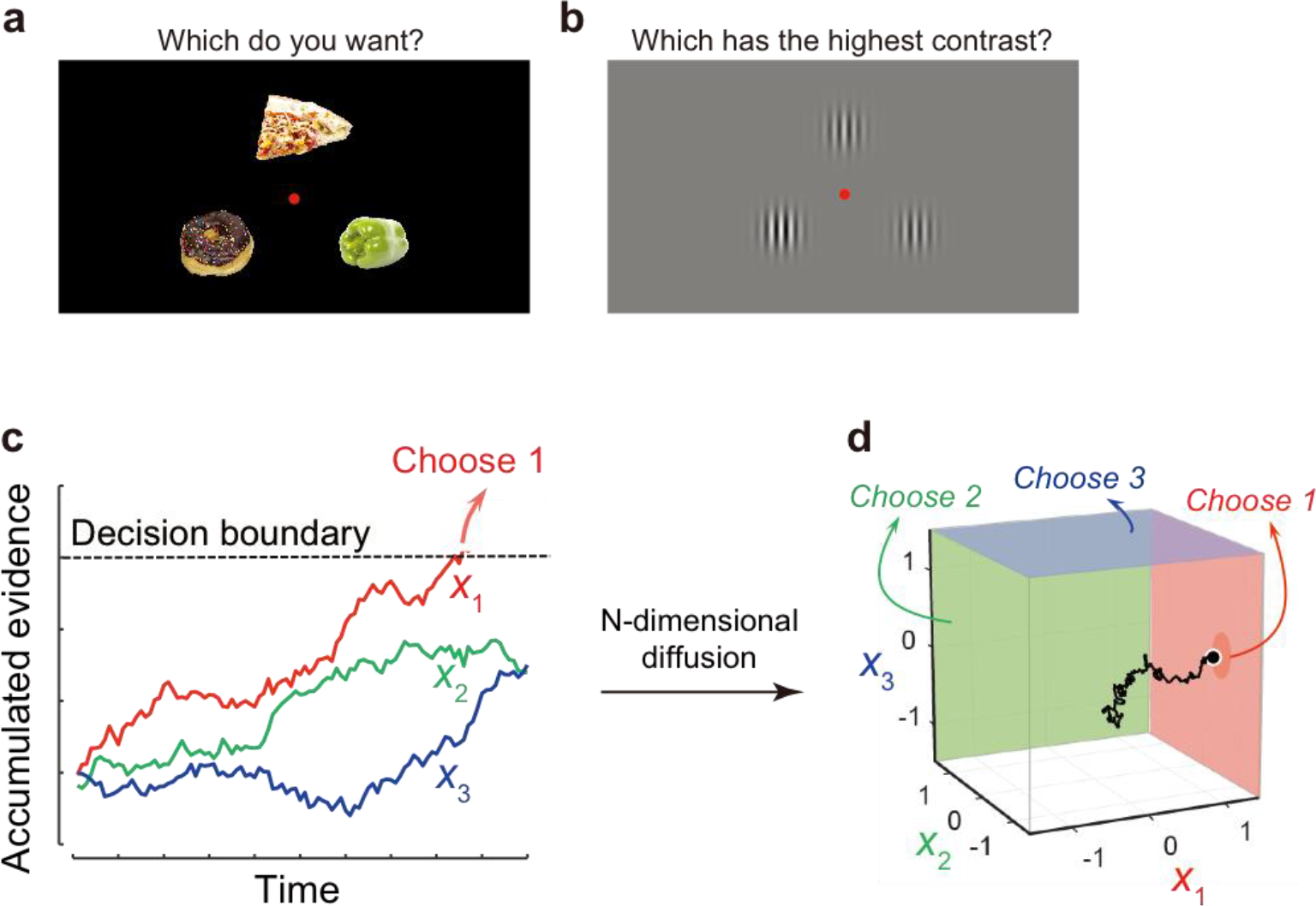
Multi-alternative decision tasks and the standard race model. (**a**) An example value-based task in laboratory settings. In a typical experiment, participants are rewarded with one of the objects they chose (in a randomly selected trial from the whole trial sequence). (**b**) An example perceptual task, in which the participants are required to choose the highest contrast Gabor patch – in this example the bottom-left one. (**c**) The race model (RM). The colored traces represent the accumulated evidence for individual options (*x*_1_, *x*_2_ and *x*_3_). In the RM, the accumulation process is terminated when either race reaches a constant decision boundary. (**d**) An alternative representation for the same RM, in which the races of accumulated evidence are shown as an *N*-dimensional diffusion. With this representation, the decision boundary for each option corresponds to a side of an *N*-dimensional cube, reflecting the independence of decision boundaries across options in the RM.

At the behavioral level, subjects choosing between 3 or more options exhibit several seemingly suboptimal behaviors, such as the similarity effect or violations of both the regularity principle and the independence of irrelevant alternatives (IIA) principle. The last effect, the violation of the IIA, refers to the observation that when choosing between two similar options, the subject’s ability to choose is affected by the presence of a third option even if this third option is never picked^11^. However, before concluding that such behaviors are suboptimal, it is critical to first derive the optimal policy and check whether it is compatible with this policy.

Here, we adopt such a normative approach. By contrast to previous models motivated by biological implementations^12–15^, we start by deriving the optimal, reward-maximizing strategy for multi-alternative decision-making, and then ask how this strategy can be implemented by biologically plausible mechanisms. To do so, we first extend a recently developed theory of value-based decision-making with binary options^7^ to general *N*-alternative cases, revealing nonlinear and time-dependent decision-boundaries in a high-dimensional belief space. Next, we show that geometric symmetries allow reducing the optimal strategy to a simple neural mechanism. This yields a novel extension of race models with time-dependent activity-normalization controlled by an urgency signal^10^.

The model provides a new perspective on how normalization and an urgency signal cooperate to implement close-to-optimal decisions for multi-alternative choices. We also demonstrate that the optimal policy is compatible with divisive normalization, a form of normalization that has been widely reported throughout the nervous system^8,9^. With this addition, and in the presence of internal variability, we report that the network replicates the similarity effect as well as violates both the independent irrelevant alternative principle and the regularity principle. Thus, our model isolates the functional components required for optimal decision-making and replicates a range of essential physiological and behavioral phenomena observed for multi-alternative decisions.

## Results

### The optimal policy for multi-alternative decisions

Suppose we have *N* alternatives to choose from in perceptual or value-based decisions. The decision maker’s aim is to make choices whose outcome depends on a-priori unknown variables (e.g., the true rewards, Figure 1a, or stimulus contrasts, Figure 1b) associated with the individual options and whose values vary across choice trials. We will assume that during the course of a decision on a given trial, each short time duration *δt* yields a piece of noisy momentary evidence about the true values of the hidden variables. For instance, in the case of perceptual decision making, this would correspond to observing new sensory information, while for value-based decision making this might be the results of recalling past experiences from memory^16^. Our derivation shows that the optimal way of accumulating such evidence is to simply sum it up over time (see **Methods)**. This reduces the process of forming a belief about these variables to a diffusion (or random walk) process, ***x***(*t*), in an *N*-dimensional space, as implemented by RMs (the black trace in Figure 1d).

Next, we derive the optimal stopping strategy: when should the decision maker stop accumulating evidence and trigger a choice? To do so, and in contrast to experiments in which subjects have to wait until the end of the trial to respond, we only consider the more natural scenario in which the decision maker is in control of their decision time. In a standard RM, evidence accumulation stops whenever one of the races reaches a threshold that is constant over time and identical across races. In other words, the evidence accumulation stops once the diffusing particle hits any sides of an *N*-dimensional (half-)cube (Figure 1d). While simple, this stopping policy is not necessarily optimal. To find the optimal policy, we utilize tools from dynamic programming^7,17,18^. One such tool is the “value function” *V*(*t*, ***x***), which corresponds to the expected reward for being in state ***x*** at time *t*, assuming that the optimal policy is followed from there on. This value function can be computed recursively through Bellman’s equation^17^. For the simple case of a single, isolated choice, the decision maker aims to maximize the expected reward (or reward rate per unit time) for this choice minus some cost *c* for accumulating evidence per unit time. One can imagine several different types of costs, such as, for example, the metabolic cost of accumulating more evidence. Once we embed this single choice within a long sequence of similar choices, an additional cost *ρ* emerges that reflects missing out on rewards that future choices yield (**Methods**). Overall, the optimal decision policy results in

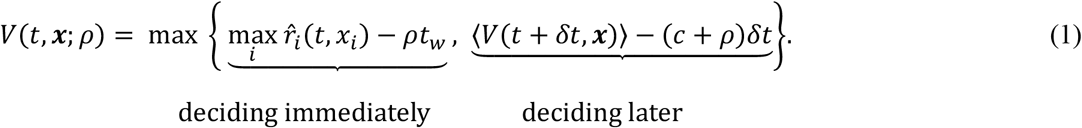

This value function compares the value for deciding immediately, yielding the highest of the *N* expected rewards 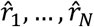, with that for accumulating more evidence and deciding later. *ρ* is the reward rate (the average reward obtained per unit time; see **Methods** for the formal definition); *t*_*w*_ is the inter-trial interval including the non-decision time required for motor movement. The expected reward for each option, 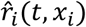 is computed by combining the accumulated evidence with the prior knowledge about the reward mean and variance through Bayes rule (**Methods**). As shown by dynamic programming theory ^19^, the larger of these two terms yields the optimal value function, and their intersection determines the decision boundaries for stopping the evidence accumulation, and thus the optimal policy. In realistic setups, decision makers make a sequence of choices, in which case the aim of maximizing the total reward becomes equivalent (assuming a very long sequence of choice) to maximizing their reward rate, which is the expected reward for either choice divided by the expected time between consecutive choices. The value function for this case is the same as that for the single-trial choice, except for that the both values for deciding immediately and for accumulating more evidence include the opportunity cost of missing out on future rewards (**Methods**).

We found the optimal policy for this general problem by computing the value function numerically^20^ from which we derived the decision boundaries (see Figure 2a). The resulting optimal decision boundaries are complex and nonlinear (Figure 2a visualizes the optimal decision boundaries, represented as 2-dimensional surfaces, for the value-based decision with *N* = 3). Clearly, the structure of the optimal decision boundaries differs substantially from that of standard RMs (Figure 1d). Importantly, we found that they have an important symmetry: they are parallel to the diagonal—the line connecting (0,0,0) and (1,1,1) (**Supplementary Note 1** shows this more formally). This symmetry implies that any diffusion parallel to the diagonal line is irrelevant to the final decision, such that we only need to consider the projection of the diffusion process onto the hyperplane orthogonal to this line (Figure 2b). The decision boundaries remain nonlinear even in this projection, as depicted by the curvatures of the solid lines in Figure 2b. Note that the nonlinearity of the decision boundaries is specific to multi-alternative choice situations (i.e., *N* ≥ 3). Indeed, for binary choices, our derivation indicates that the projection of the diffusion process onto an *N* − 1 dimensional subspace becomes a projection onto a line since *N* = 2. On this line, the stopping boundaries are just two points and therefore cannot exhibit any nonlinearities. In fact, for *N* = 2, the optimal policy corresponds to the well-known drift diffusion model of decision making^7,17^.

**Figure 2.**
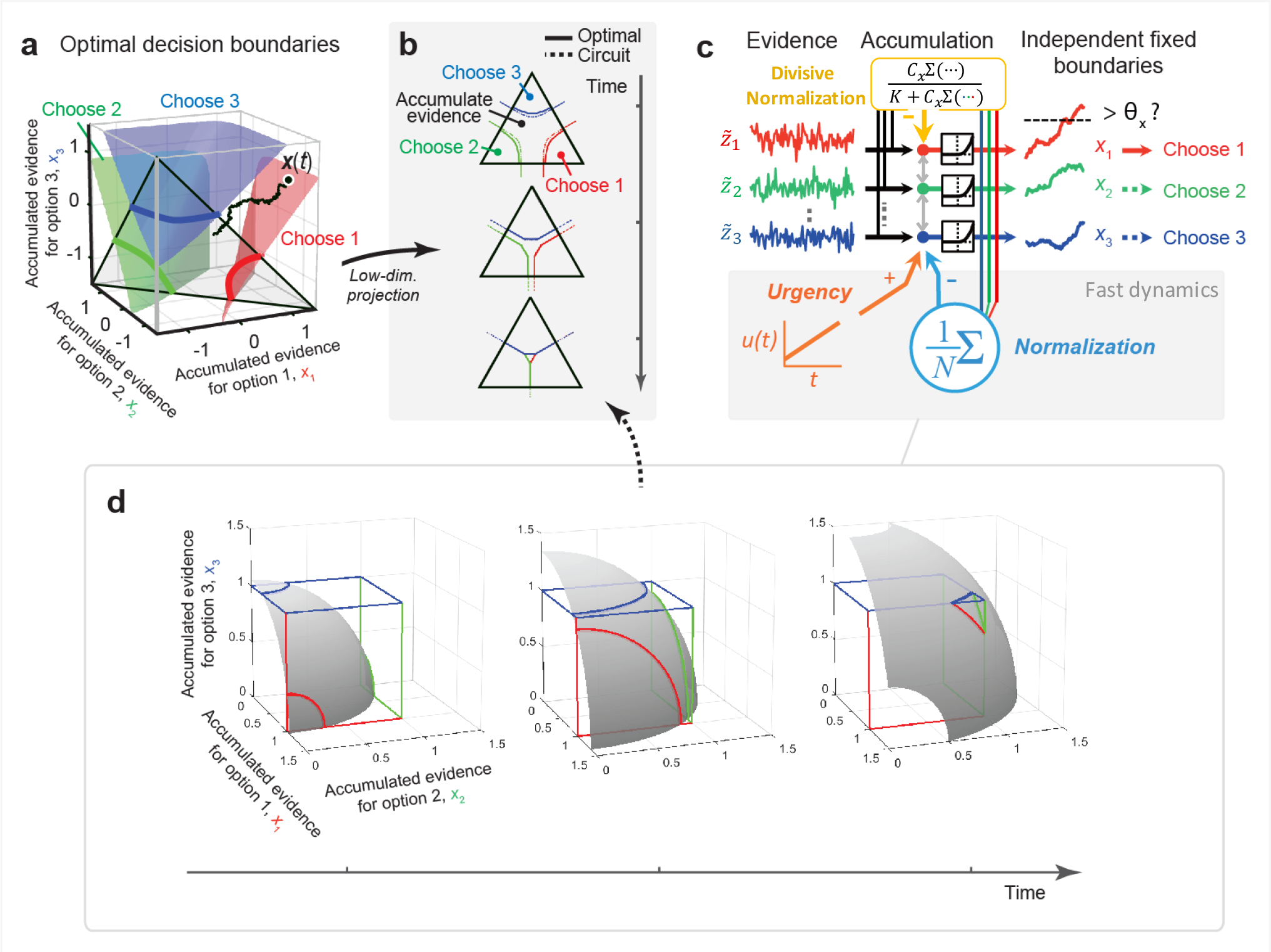
The optimal decision policy for 3-alternative choices. (**a**) The derived optimal decision boundaries in the diffusion space. In contrast to the standard race model’s decision boundaries (Figure 1d), they are nonlinear, but symmetric with respect to the diagonal (i.e., the vector (1,1,1)). (**b**) Lower-dimensional projections of decision boundaries at different time points. The solid curves are the optimal decision boundaries projected onto the plane orthogonal to the diagonal (the black triangle in panel **a**). The dashed curves indicate the effective decision boundaries implemented by the circuit in panel **c**. (**c**) The circuit approximating the optimal policy. Like RMs, it features constant decision thresholds that are independently applied to individual options. However, the evidence accumulation process is now modulated by recurrent global inhibition after a nonlinear activation function (the “normalization” term) and a time-dependent global bias input (“urgency signal”), and rescaled (“divisive normalization”). (**d**) Schematic illustrations of why the circuit in panel **c** can implement the optimal decision policy. The nonlinear recurrent normalization and urgency signal constrain the neural population states to a time-dependent manifold (the gray surfaces). Evidence accumulation corresponds to a diffusion process on this nonlinear (*N* − 1 dimensional) manifold. The stopping bounds are implemented as the intersections (the colored thick curves) of the manifold and the cube (colored thin lines), in which the cube represents the independent, constant decision thresholds for the individual choice options. Divisive normalization rescales the space of evidence accumulation, leaving the relative distances between the accumulators and stopping bounds intact (not shown). Due to the urgency signal, the manifold moves toward the corner of the cube as the time elapses, causing the intersections (i.e., the stopping bounds) to collapse onto each other over time.

Numerical solutions also revealed that the optimal decision boundaries evolve over time: they approach each other as time elapses, and finally collapse (Figure 2b, solid curves). These nonlinear collapsing boundaries differ from the linear and static ones of previous approximate models, such as multi-hypothesis sequential probability ratio tests (MSPRTs)^21–23^, which are known to be only asymptotically optimal under specific assumptions (**Methods**).

We show in the **Supplementary Note 4** that these results generalize to models in which the streams of noisy momentary evidence are correlated in time, either with short range temporal correlations, as is often observed in spikes trains, or with long range temporal correlations as postulated for instance in the linear ballistic accumulator model^24,25^. Our results also apply to experiments such as the ones performed by Thura and Cisek^26,27^ in which the momentary evidence are accumulated directly on the screen, in which case there is no need for latent integration but the stopping bounds on the observed accumulated evidence remain the same as in Figure 2a.

### Circuit implementation of the optimal policy

In the optimal policy we have derived, the evidence accumulation is simple: it involves *N* accumulators, each of which sum up their associated momentary evidence independent of the other accumulators. By contrast, the stopping rule is complex: at every time step, the policy requires computing *N* time-dependent nonlinear functions that form the individual stopping boundaries. This rule is nonlocal because whether an accumulator stops depends not only on its own state but also on that of all the other accumulators. A simpler stopping rule would be one in which a decision is made whenever one of the accumulators reaches a particular threshold value, as in independent RMs. This, however, would require a nonlinear and nonlocal accumulation process in order to implement the same policy through a proper variable transformation. Nonetheless, such a solution would be appealing from a neural point of view as it could be implemented in a nonlinear recurrent network endowed with a simple winner-take-all mechanism that selects a choice once the threshold is reached by one of the accumulators.

Armed with this insight, we found that a recurrent network with independent thresholds, as depicted in Figure 2c, can indeed approximate the optimal solution very closely. It consists of *N* neurons (or *N* groups of identical neurons), one per option, which receive evidence for their associated option. The network operates at two time-scales. On the slower time-scale, neurons accumulate momentary evidence independently across options according to:

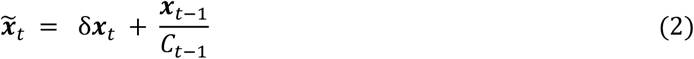

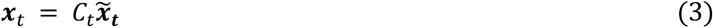

where ***x***_*t*_ is the vector of accumulated evidence at time *t*, *δ**x***_*t*_ is the vector of momentary evidence at time *t* and *C*_*t*_ is the commonly used divisive normalization,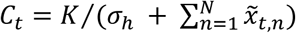, (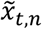 denotes the *n*^th^ component of the vector 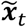 at time t) which is found all over cortex and in particular in LIP^8^.

On the faster time scale, activity is projected onto a manifold defined by 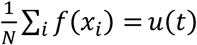, (shown as a gray surface in Figure 2d) where *u(t)* is the urgency signal. This operation is implemented by iterating:

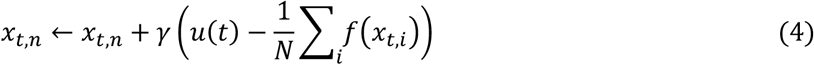

until convergence, *γ* is the update rate, and *f* is a rectified polynomial non-linearity (see **Methods** and **Supplementary Note 2** for further details). This process is stopped whenever one of the integrators reaches a preset threshold. The choice of this projection was motivated by two key factors. First, this particular form ensures that the projection is parallel to the diagonal, the line connecting (0,0,0) and (1,1,1). As we have seen, diffusion along this axis is indeed irrelevant. Second, the use of a nonlinear function *f* implies that we do not merely project on the hyperplane orthogonal to the diagonal. Instead, we project onto a nonlinear manifold. This step is what allow us to approximate the original complex stopping surfaces with simpler independent bounds on each of the integrators as illustrated in Figure 2d (see **Supplementary Note 2** for a formal explanation). The time dependent urgency signal, *u(t)*, implements a collapsing bound, which is also part of the optimal policy (Figure 2b). Indeed, this urgency signals brings all the neurons closer to their threshold and, as such, is equivalent to the collapse of the stopping bounds over time (Figure 2d).

Equations 2, 3, and 4 can be turned into a single differential equation (see Eq. 40 in **Supplementary Note**). The iterative difference equations we show here are a particular form of the implementation, making it easier to interpret the diffusion process. Importantly, Equations 2 and 3 provide a generalization of divisive normalization which ensures that evidence is still integrated optimally over time.

The model contains three parameters: the power of the nonlinearity, and the starting point and slope of the urgency signal (**Methods**). When these parameters are optimized to maximize reward rate, the network approximates very closely the optimal stopping bounds (Figure 2b). As a result, the reward rate achieved by the network is within 98% and 95% of the optimal reward rate for 3 and 4 options, respectively (across a wide range of prior distributions over rewards, **Methods**).

A simple extension of this network can be used to model other types of task such as the one used by Thura and Cisek ^26,27^ in which the momentary evidence available on the screen is in fact the accumulated evidence since the beginning of the trial. All that is needed is to remove the temporal integration step in the input neuron, since this step is now superfluous.

### Normalization and urgency improve the task performances

Our circuit model comprises independent decision thresholds for individual options, as in standard RMs (consistent with recordings in the parietal cortex ^10^), but features time-dependent normalization in addition to an urgency signal. To quantify the contribution of each circuit component, we compared the performance of four different circuit models: (i) the standard RM with independent evidence accumulation within each accumulator, (ii) an RM with the urgency signal alone, (iii) an RM with normalization alone, and (iv) the full model with both the urgency signal and normalization.

This comparison revealed that adding the urgency signal and/or normalization to the standard RM indeed improved the reward rate. Intriguingly, normalization had a much larger impact than the urgency signal (Figure 3, **left**), demonstrating the relative importance of normalization in improving the reward rate. The performance differences across models shrink with an increasing number of options because performance shown here is relative to a model making random, immediate choices. Indeed, as the number of options to choose from increases, the absolute reward rate of the full and reduced models increases at similar rates, while the performance of the random model remains the same because, for value-based decisions, this policy simply effectively draws a random sample from the prior over rewards on each trial regardless of how many choices are available. As a result, the performance of the different models relative to the random model (as shown in Figure 3) become more similar.

**Figure 3.**
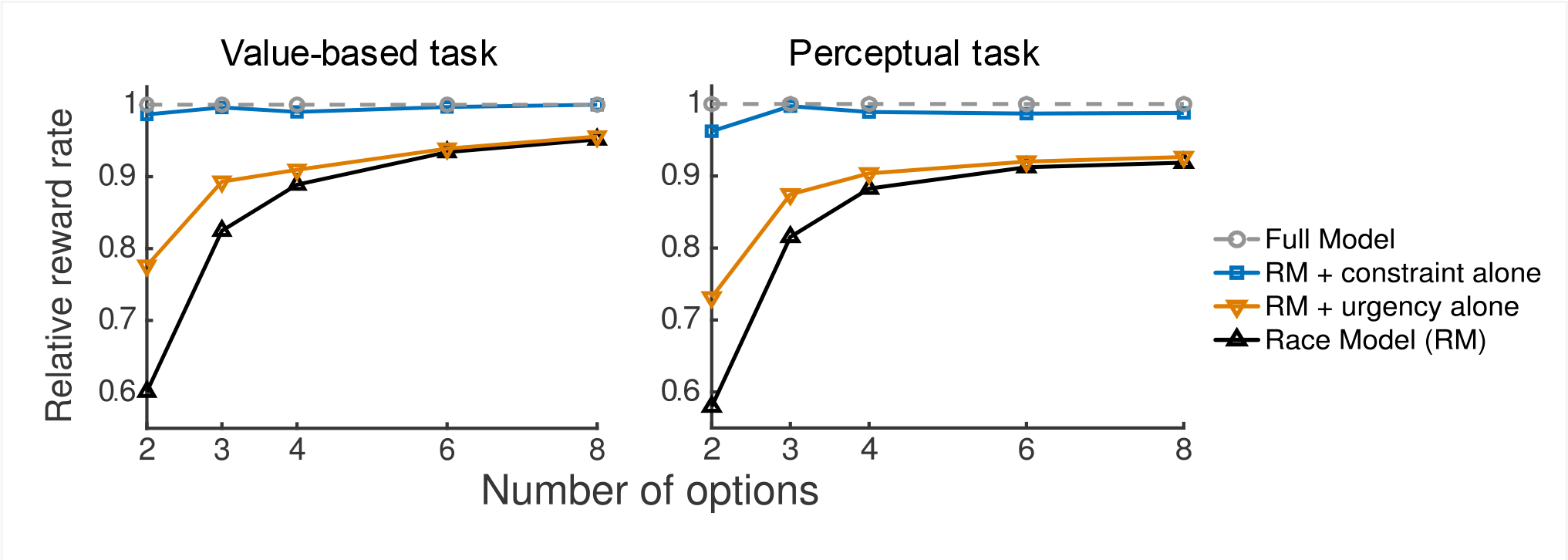
Normalization and urgency improve the task performance. Relative reward rates in value-based (left) and perceptual tasks (right). To quantify the contribution of each circuit component, we compared the performance of four different circuit models: (i) the standard race model (RM) with independent evidence accumulation within each accumulator, (ii) an RM with only an urgency signal, (iii) an RM with only normalization, and (iv) the full model with both urgency signal and normalization. We quantified the reward rates of models 1-3 (“ reduced models”) relative to that of the full model by 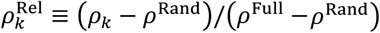, where *ρ*_*k*_(*k* = 1,2,3) denotes the reward rates of reduced models 1–3; 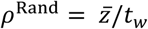 is the baseline reward rate of a decision-maker who makes immediate random choices after trial onset. *ρ*^Full^ is the reward rate of the full model with both normalization and urgency.

The overall results were similar for perceptual decisions, in which the decision-maker is rewarded based on whether the response is correct or incorrect (Figure 3, **right**). Thus, normalization and urgency signal contribute to improving task performance in both value-based and perceptual decision-making tasks.

### Relation to physiological and behavioral findings

#### Urgency signal

We examined how neural dynamics and behavior predicted by the proposed circuit relates to previous physiological and behavioral findings. First, we found that the average activity in model neurons rises over time, independently of the sensory evidence, consistent with the urgency signals demonstrated in physiological recordings of neurons in the lateral intraparietal cortex (LIP)^10^ (Figure 4a). Interestingly, our model also replicates a gradual decrease in the slope of the average neural activity over time which arises in the model as a consequence of the nonlinear recurrent process. In typical physiological experiments, urgency signals are extracted by averaging over neural activities across the entire recorded population, including different stimulus conditions. The rationale behind this procedure is that the urgency signal has been postulated as a uniform additional input to all parietal neurons involved in the evidence accumulation process. A signal extracted this way differs from the function *u*(*t*) in our model, which is followed by constraining the activity nonlinearly through recurrent neural dynamics, and thus does not trivially relate to the empirically observed urgency signals. Nonetheless, the average activity in model neurons was, through simulations, found to replicate the temporal increase, consistent with the physiological recording in LIP neurons ^10^ (Figure 4b).

**Figure 4.**
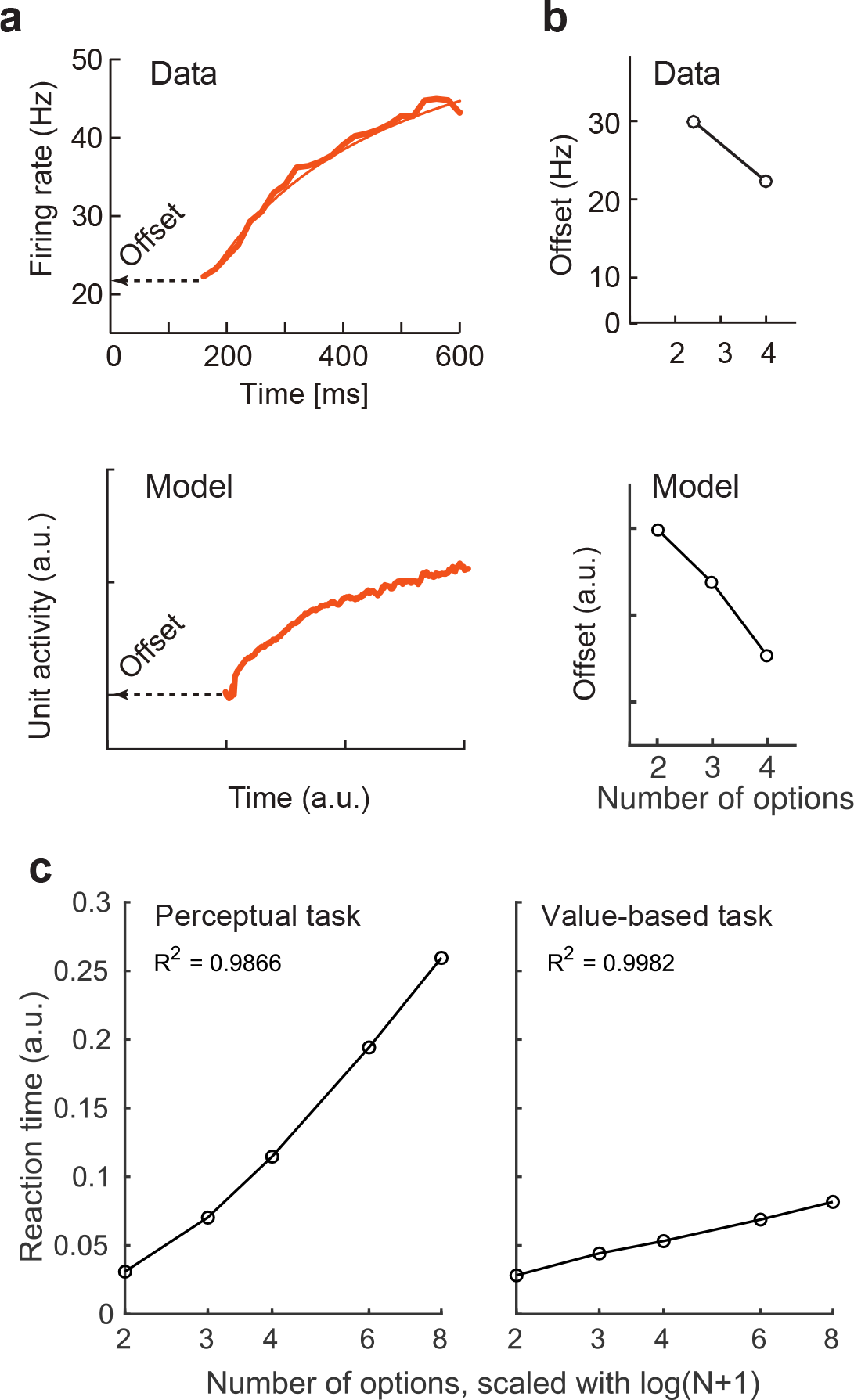
The model replicates the neuronal urgency signal and Hick’ law in choice reaction times (RTs). (**a**) Urgency signals in lateral intraparietal (LIP) cortex neurons (top) and in the model (bottom). In typical physiological experiments, urgency signals are extracted by averaging over neural activities across the entire recorded population, including different stimulus conditions. The rationale behind this procedure is that the urgency signal has been considered as a uniform additional input to all parietal neurons involved in the evidence accumulation process. A signal extracted this way is not exactly the same as the global input signal (function *u*(*t*) in Figure 2c) to the circuit, which includes nonlinear activity normalization through recurrent neural dynamics, and thus does not trivially relate to the empirically observed urgency signals. Nonetheless, the average activity in model neurons was found to replicate the temporal increase, including the saturating temporal dynamics. (**b**) The initial offset activities decrease with an increasing number of options, in both LIP neurons (top) and the model (bottom). The data figure was modified from Ref. ^10^. (**c**) The choice RTs following Hick’s law. The RTs increase with the number of options (*N*) in both perceptual (left) and value-based (right) tasks. Note the logarithmic scaling of the horizontal axis.

#### Decrease in offset activities in multi-alternative tasks

Second, it has been reported that the initial “offset” (i.e., the average neural activity) of evidence accumulation^10,28^ decreases as the number of options increases (Figure 4b), although to our knowledge no normative explanation has been offered for this observation. Interestingly, our circuit model replicates this property when optimized to maximize the reward rate (Figure 4b). For a fixed number of options, and for the particular type of sequential decisions we are considering here, lowering the initial offset increases both accuracy and reaction time (RT), but has a proportionally stronger effect on accuracy such that the reward rate increases. On the other hand, increasing the number of options while leaving the initial offset unchanged causes a decrease in both accuracy and reaction time, and an associated drop in reward rate. Thus, to counter-act this drop, we need to lower the initial offset, resulting in a lower optimal offset for a larger number of options. The change in the optimal offset size also explains the behavioral effects in RTs as described below.

#### Hick’s law in choice RTs

Third, the change in the optimal offset also explains the behavioral effects in choice RTs known as “Hick’s law”^29,30^. Hick’s law is one of the most robust properties of choice RTs in perceptual decision tasks^29,30^. In its classic form, it suggests the linear relationship *RT* = *a* + *b*log(*N* + 1) between mean RT and the logarithm of the number of options (*N*). Our model replicates this near-logarithmic relationship (Figure 4c). The increased RT for a larger number of options is concordant with the decrease in offset activities as described in the previous section. Interestingly, the RT dependency on the number of options tends to be much weaker for value-based than perceptual decisions.

#### Value normalization

Fourth, our model replicates suppressive effects of neurally encoded values among individual options. In particular, the activity of the LIP neurons encodes values of targets inside the neuronal receptive fields, but is also affected by values associated with targets displayed outside the receptive fields^8,9,31^. The larger the total target values outside these receptive fields, the lower the neural activity, which is usually described as normalization. The model replicates these suppressive effects (Figure 5a).

**Figure 5.**
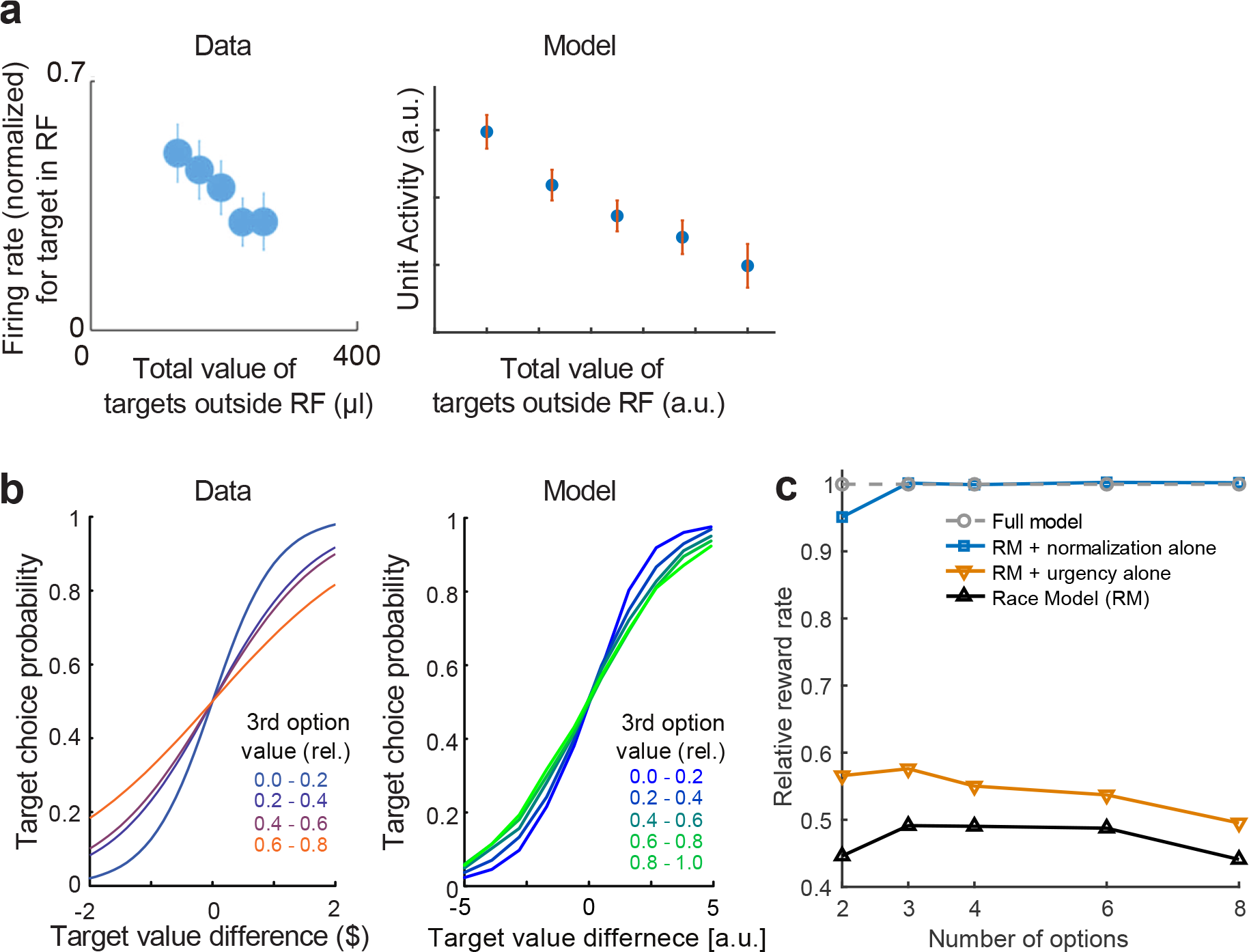
Activity normalization and violation of the axiom of independence of irrelevant alternatives (IIA). (**a**) Neuronal response to a saccadic target associated with a fixed reward as a function of the total amount of reward for all other targets on the screen in the lateral intraparietal area (left) and in the model (right). The data figure was modified from Ref ^8^. In both LIP and the model, the response of a neuron to a target associated with a fixed amount of reward decreases as the reward to the other targets increases. In the model, this effect is induced by the normalization. (**b**) Left plot: as the value of a third option is increased, the psychometric curve (for a fixed decision time, set by the experimenter) corresponding to the choice between options 1 and 2 becomes shallower – a result that violates the axiom of IIA. Data figure was modified from Ref. ^11^. Right plot: the model with added neural noise after activity normalization exhibits the same behavior. (**c**) In the presence of internal variability, the race model variants without constrained evidence accumulation approximating the optimal policy (second term in Equation 2) perform much worse than our model variants with that constraint (when compared to Figure 2d).

#### Violation of IIA

So far, our neural model only has one source of variability, namely, the noise corrupting the momentary evidence. There are, however, other sources of variability that are quite likely to exist in brain. For instance, the decision maker must learn how to properly adjust the decision bounds in order to optimize reward rate, which would result in variability in the value of the bound from trial to trial. There is experimental evidence suggesting that learning can indeed induce extra variability in decision making tasks^32^. The variability in these bounds could also be purposely induced by neural circuits to ensure that the decision maker does not always choose the option with the highest value but also explore alternatives. Such an exploration behavior is critical in environments in which the value of the options varies over time, which is common in real world situations.

In our neural model, we added such extra variability directly to the accumulator, which is mathematically equivalent to adding it to the bound. Despite this extra variability, our neural model continues to outperform the race model (Figure 5c). Stripping the normalization from the full model results in a large drop in reward rate with a further drop, though less pronounced, when the urgency signal is also removed.

Importantly, this version of the model also replicates apparently “irrational” behavior in humans and animals which violates the principle of “independence of irrelevant alternatives (IIA)”^33^, an axiomatic property assumed in traditional rational theories of choice^34,35^. Behavioral studies have shown that the choice between two high-valued options depends on the value of a third alternative option^36–41^, even if the value of this third option is so low that it is never chosen. One example of such an interaction is shown in Figure 5b. In this experiment, subjects found it increasingly harder to pick among their two top choices as the value of the third option is increased. Our noisy neural model exhibits a similar violation of the IIA (Figure 5b), which is primarily caused by the divisive normalization. The divisive normalization decreases the mean value difference between the two top options as the value of the third option is increased, making these two options harder to distinguish due the presence of internal variability.

#### Violation of the regularity principle

In multi-alternative decision making, subjects not only violate the IIA but also the regularity principle. The regularity principle asserts that adding extra options cannot increase the probability of selecting an existing option. We have found that the same model that violates the IIA also violates this regularity principle. At first, this may seem counterintuitive. Introducing a third option into a choice set must decrease the probability of picking either of the first two options, which is consistent with the regularity principle. However, consider the probability of picking option 1 when option 2 is more valuable. In the absence of a third option, this probability will tend to be very small. When the third option is introduced, and its value is increased, the violation of the IIA implies that the probability of picking option 1, relative to option 2, will increase, as illustrated by the shallower psychometric curves in Figure 5b. Therefore, two factors are at play here, with opposite effects: the presence of a third option implies that choices 1 and 2 are picked less often, but the probability of picking option 1 increases as a result of the IIA violation. Our simulations reveal that the second factor dominates when the value of option 1 is smaller than that of option 2, as illustrated in Figure 6a.

**Figure 6.**
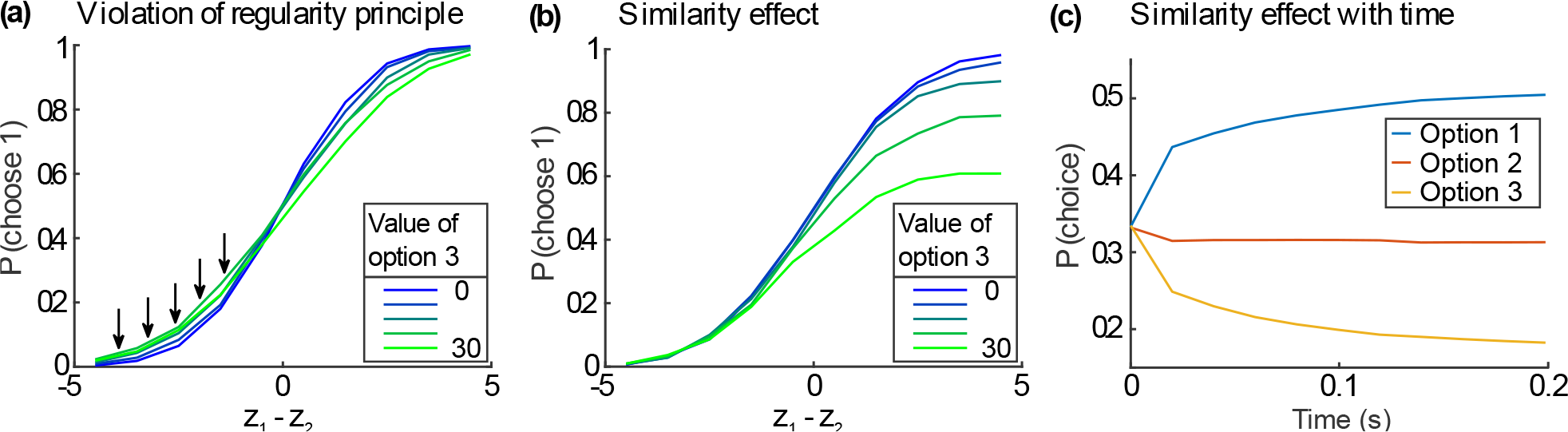
Regularity and similarity principles. (**a**) Violation of the regularity principle. When a third choice is introduced, the probability of choosing option 1 increases as the value of option 3 increases. This effect is only observed option 1 is much less valuable than option 2. (**b**) The similarity effect: Adding a third option, similar to option 1, reduces the probability of choosing option 1 relative to option 2 as the value of option 3 increases. The inset shows that the probability of picking option 1 also decreases as the value of option 3 increases. (**c**) The strength of the similarity effect increases with time within the course of a single trial, as shown by the decrease in the probability of choosing option 1 as time elapses.

#### The similarity effect

Our model also replicates the similarity effect that has been reported in the literature^40, 42, 43^. This effect refers to the fact that when subjects are given a third option which is similar to, say, option 1, the probability of choosing option 1 decreases. To model this effect, we postulated that each object is defined by a set of features and that its overall value is a linear combination of the values of its features. As before, we also assume that the values of the features are not known exactly. Instead, the brain generates noisy samples of these values over time. In this scenario, the similarity between two objects is proportional to the overlap between their features. This overlap implies that the stream of value samples for the two similar options are correlated, while being independent for the third, dissimilar option. Accordingly, we simulated a 3-way race in which the momentary evidence for options 1 and 3 are positively correlated. As illustrated in Figure 6b, we found that the probability of choosing option 1 decreases relative to option 2 as the value of option 3 increases, thus replicating the similarity effect. As has been observed experimentally ^44,45^, we found that the similarity effect grows over time during the course of a single trial Figure 6c.

### Predictions

Our model makes a number of experimental predictions at both the behavioral and neural levels (see **Supplementary Note 3** for further details).

First, during evidence accumulation, the neural population activity should be near an *N* − 1 dimensional continuous manifold (i.e., a nonlinear surface), where *N* is the number of choices (Figure 2d). This is a direct consequence of evidence accumulation paired with nonlinear normalization. As the activity of *D* neurons is *D*-dimensional, and since *N* ≪ *D* in general, our prediction implies that neural activity should be constrained to a small subspace of the neural activity space. This prediction can be tested with standard dimensionality reduction techniques using multi-electrode recordings although this analysis should be done carefully since our model also predicts that the position of this manifold changes over time. Failure to take this time dependency into account could significantly bias the estimate of the dimensionality of the constraining manifold. Our theory makes 11 additional predictions related to existence and properties of the manifold which are listed in **Supplementary Note 3**.

Second, our model correctly predicted the decrease in the initial activity offset value of LIP neurons with the number of choices (the offset is the baseline firing rate value right before evidence accumulation). Remarkably, this offset decrease results from an economic strategy that maximizes the reward rates by balancing the speed and accuracy in a long sequence of trials under the opportunity cost for future rewards. Thus, the offset should also be modulated by other reward rate manipulations. In particular, we predict that increasing the average reward rate by either increasing the reward associated with the choices or decreasing the inter-trial interval should raise the offset for a fixed number of choices.

Third, previous studies have considered two types of strategies for multi-alternative decision making: the ‘max-vs.-average’ and the ‘max-vs.-next’^6,46,47^. In the former, the winning race is the first one to reach a particular difference between its own state and the average of the other races (Figure 7b). In the ‘max-vs.-next’, it is the difference between the top race and the second best one that matters (Figure 7c). Our theory predicts that subjects should smoothly transition between these two modes depending on the pattern of rewards across choices (Figure 7a), a prediction that can be tested with standard psychophysics experiments. If all choices are equally rewarded, our model predicts that subjects should adopt a ‘max-vs.-average’ strategy. This is because the particle, which starts from the center of the triangle in Figure 7a, will tend to diffuse equally well in all directions and will therefore hit the optimal bounds close to where they overlap with the bounds corresponding to the ‘max-vs.-next’ strategy (Figure 7b, d). If only two choices are highly rewarded, our model switches to a ‘max-vs.-next’ strategy because the particle will quickly drift toward the side of the triangle corresponding to the two high valued choices where the optimal bounds overlap most with the bounds corresponding to the ‘max-vs.-average’ strategy (Figure 7c,d). If only one option is highly rewarded, our model reverts to the ‘max-vs.-average’ model (Figure 7b,e). Therefore, our model predicts that if humans follow the optimal strategy, they should show similar transitions between the ‘max-vs.-average’ and the ‘max-vs.-next’ strategies.

**Figure 7.**
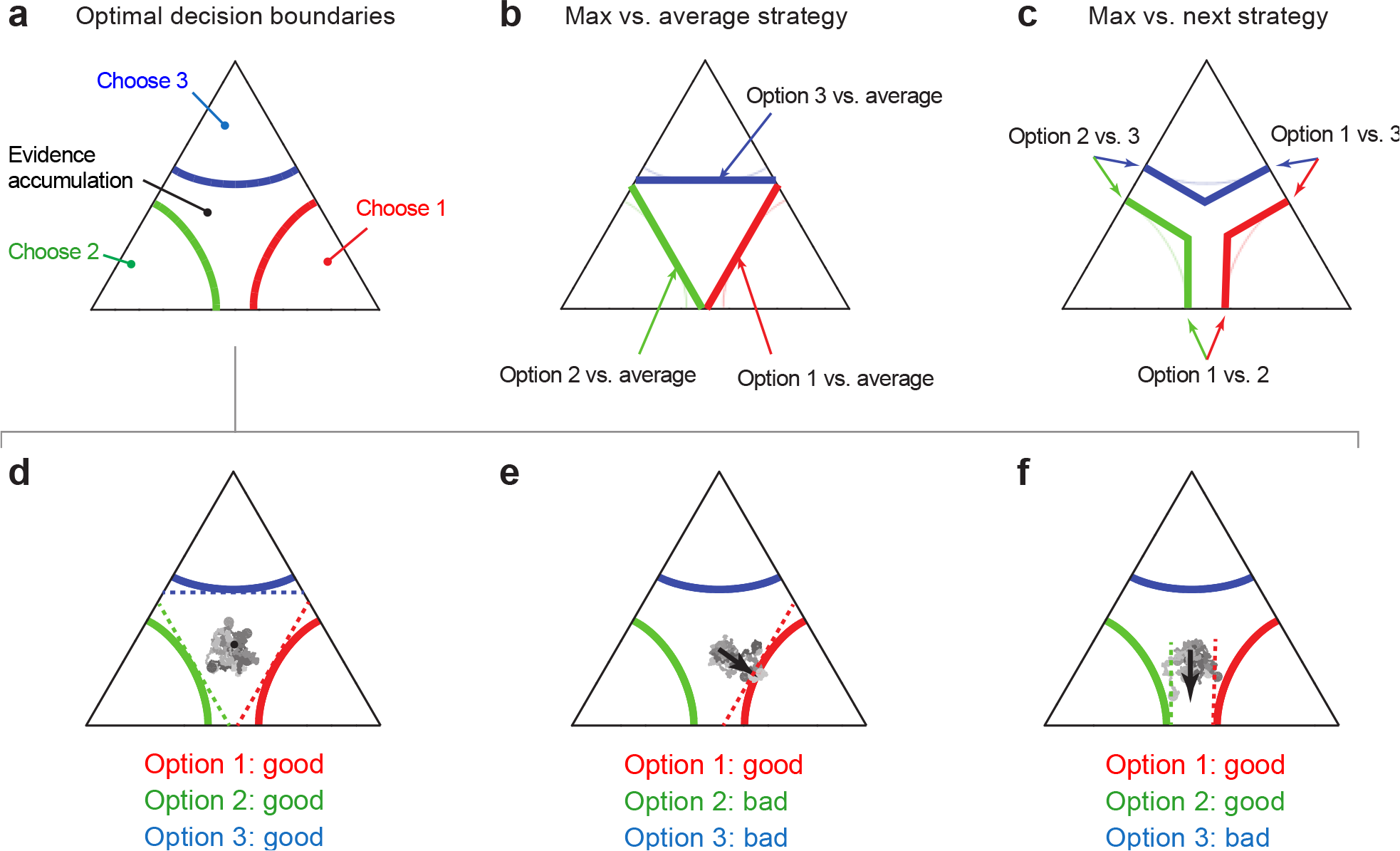
The optimal policy predicts a smooth transition between the ‘max vs. next’ and the ‘max vs. average’ decision strategies depending on the relative values of the three options. (**a**) The stopping bounds for the optimal policy after projecting the diffusion onto the hyperplane orthogonal to the diagonal. (**b**) The stopping bounds corresponding to the ‘max vs. average” strategy (thick colored lines). In this strategy, the decision-maker computes the difference between each option’s value estimate and the average of the remaining options’ values and triggers a choice when this difference hits a threshold. The stopping bounds in this case overlap with the optimal bounds from panel **a** (shown here as thin colored lines) in the center but not on the side. (**c**) The stopping bounds for the ‘max vs. next’ strategy (thick lines). In this strategy, the decision-maker compares the best and second-best value estimates, and makes a choice when this difference exceeds a threshold. In a three-alternative choice, this is implemented with three pairs of linear decision boundaries (colored thick lines) corresponding to the three possible combinations of two options. In contrast to the bounds for the max vs average strategy, the bounds for the max vs next strategy overlap with the optimal bounds (thin colored lines) on the edge of the triangle but not in the center. (**d**) When all three options are equally good, the diffusion of the particle is isotropic and is therefore more likely to hit the stopping bounds in their centers, where they overlap with the max vs average strategy. (**e**) When one option is much better than the other two, the diffusion is now biased toward the center of the bound corresponding to the good option, which is once again equivalent to the max vs average strategy. (**f**) When two options are equally good, while the third is much worse, the particle will tend to drift toward the part of the triangle corresponding to the two good options (black arrow), where the optimal bound overlaps with the bounds for the max vs next strategy. The grey curves in **d**-**f** illustrate accumulator trajectories that are typical for the considered scenarios.

## Discussion

In this study we discussed the optimal policy for decisions among more than two valuable options, as well as a possible biological implementation. The resulting policy has nonlinear boundaries and thus differs qualitatively from the simple diffusion models that implement the optimal policy for the two-alternative case^7^. More specifically, this work makes four major contributions. First, we prove analytically that the optimal policy involves a nonlinear projection onto an *N* − 1 dimensional manifold, which can be closely approximated by neural circuits with a nonlinear normalization. Second, apparently “irrational” choice behaviors, such as the violation of the IIA, are reproduced by our optimal model in the presence of variability arising, for instance, from learning or exploration. Third, we found that the distance to threshold must increase with set size for optimal performance. This has already been observed experimentally^10,28^ (Figure 4a), but no computational explanation has ever been offered for this effect until now. Fourth, the model follows Hick’s law, that is, it predicts that reaction times in value-based decisions should be proportional to the log of the number of choices plus one, as is commonly observed in behavioral choice data. However, our model does not account for violation of Hick’s law for saccadic eye movements effects ^48,49^, or the well know pop-out effect reported in visual search, in which reaction times are independent of the number of items on the screen ^50^. Capturing these effects would require that we specialize our model to the specific context of these experiments which lie beyond the scope of the present manuscript.

Our replication of the violation of the IIA is similar to what Louie et al have recently^11^ proposed though, in their case, they did not consider noise in the momentary evidence and they did not derive the optimal policy for multi-alternative decision making. Therefore, our work is the first one to demonstrate that an optimal policy for multi-alternative decision making using divisive normalization violates the IIA in the presence of internal noise. Preliminary work by Steverson et al.^51^ has also clarified the conditions under which networks with divisive normalization implement the optimal policy for decision making with respect to internal noise, thus suggesting that divisive normalization is indeed required for optimal decision making when all sources of noise are considered. Moreover, recent proof of equivalence between divisive normalization and an information-processing model offers another explanation for the role of divisive normalization: to optimally balance the expected value of the chosen option with the entropic cost of reducing uncertainty in the choice^51^.

A well-known strategy to decide among multiple options is the MSPRT^21,22^, and previous studies have shown that the MSPRT could be implemented/approximated by neural circuits^23,47,52^. However, the MSPRT has not been designed for the problems we consider here. First, it assumes that the decision-maker receives a fixed magnitude of reward based on the accuracy of choices (i.e., whether they are correct or incorrect) in each trial, as in conventional perceptual decision tasks. Value-based decisions, in which the reward magnitude can vary across trials, clearly violate this assumption. Second, it furthermore assumes a constant task difficulty whereas the present study assumes the difficulty of both value-based and perceptual choices to vary across these choices. Third, since the MSPRT is only asymptotically optimal in the limit of infinitely small error rates (i.e. when the model’s performance is near 100% correct), it deviates from the optimal policy when this error rate is not negligible^21,22^. Our present analysis clarifies the properties of the optimal decision policy under multiple options, which differs from the MSPRT by characteristic nonlinear and collapsing decision boundaries. Despite the apparent complexity of those decision boundaries, we found that a symmetry in these boundaries allows the optimal strategies to be approximated by a circuit that features well-known neural mechanisms: RMs whose evidence accumulation process is modulated by normalization, an urgency signal, and nonlinear activation functions. The model provides a consistent explanation for the functional significance of normalization and urgency signal: they are necessary to implement optimal decision policies for multi-alternative choices in which subjects control the decision time.

Although we modeled the uncertainty about the true hidden states or values with a single Gaussian process that represents the noisy momentary evidence, in realistic situations the uncertainty could have multiple origins, including both external and internal sources. Potential sources of the external noises include the stochastic nature of stimuli, sensory noise, and incomplete knowledge about the options (e.g., having not yet read the dessert of a particular menu option when choosing among different lunch menus). On the other hand, internal noises could result from learning, exploration, uncertain memory or ongoing value inference (e.g., sequentially contemplating features of a particular menu course over time). We assumed simplified generative models with an unbiased and uncorrelated Gaussian prior; future extensions should cover more complex setups including asymmetric mean rewards among options.

Note that the present study considers the simplified case in which the value of each option is represented with a scalar variable. We have shown that this model is sufficiently complex to replicate basic behavioral properties such as Hick’s law, the violation of IIA, the similarity effect, and the violation of the regularity principle in multi-alternative choices. Future studies should cover more complex situations including value comparisons based on multiple features (e.g., speeds and designs of cars), which can lead to other forms of context-dependent choice behavior^39,40,53^. Decision-making with such a multidimensional features space requires to compute each option’s value by appropriately weighting each feature. Some studies suggest that apparently irrational human behavior could be accounted for by heuristic weighting rules for features, which integrate feature valences through feedforward^42,43,46^ or recurrent^12,44,45^ neural interactions. Interestingly, a recent study reports that a context-dependent feature weighting can increase the robustness of value encoding to neural noise in later processing stages^43,54^. However, to our knowledge, the optimal policy for these more complex models in which the value function is computed by combining multiple features, presented sequentially, remains unknown. Once this policy is derived, it will be interesting to determine whether all, or part, of the seemingly irrational behaviors that have been reported in the literature are a consequence of using the optimal policies for such decisions or genuine limitations of the human decision-making process.

Finally, the current model provides several interesting predictions on neural population dynamics: because of the normalization, the collective neural activity could be constrained to a low-dimensional manifold during decision making. The dimensionality of this manifold depends on the number of options (*N* − 1 dimensions for *N*-alternative choices) whereas the position of the manifold should depend on time, reflecting the effect of the urgency signal. These predictions could be tested with neurophysiological population recordings combined with advanced dimensionality reduction techniques.

## Author contributions

S.T., J.D. and A.P. conceived the study. S.T. and J.D. developed the theory and conducted the mathematical analysis. S.T., J.D., and N.P. performed the simulations. S.T., J.D., N.P., and A.P. interpreted the results and wrote the paper.

## Acknowledgement

A.P. was supported by the Swiss National Foundation #31003A_143707, and a grant from the Simons Foundation (#325057). J.D. was supported by a Scholar Award in Understanding Human Cognition by the James S. McDonnell Foundation.

## Methods

### Task structure and generative models

We consider *N*-alternative value-based or perceptual decisions in which the decision-maker responds as soon as she commits to a choice. Value-based and perceptual decisions differ in how choices are associated with reward: in the value-based case the decision-maker reaps the reward associated with the chosen item (e.g., a food item), whereas in perceptual paradigms the amount of reward depends only on whether the choice is “correct” in the current task contexts. In contrast to previous models motivated by biological implementations^12–15^, we start by deriving the optimal, reward-maximizing strategy for multi-alternative decision-making tasks without assuming specific biological implementations, and then ask how this strategy can be implemented by biologically plausible mechanisms. The following formulation applies to both perceptual and value-based tasks.

Let ***z*** ≡ (*z*_1_,…, *z*_*N*_) denote hidden variables (e.g., reward magnitudes for value-based tasks, or stimulus contrasts for perceptual tasks) associated with *N* choice options. These true hidden variables vary across trials, and are never observed directly and as such unknown to the decision-maker. Instead, the decision maker observes some noisy momentary evidence with mean ***z**δt*,

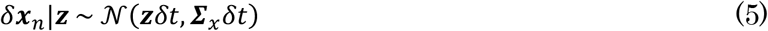

for each option *i* ∈ {1,.., *N*}, in every small time-step *n* of duration *δt*. ***∑***_*x*_ here denotes the covariance matrix of the momentary evidence. Before observing any evidence, the decision-maker is assumed to hold a normally distributed prior belief,

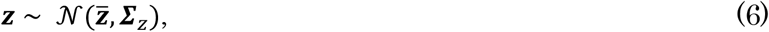

with mean 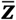 and covariance ***∑***_*z*_ reflecting the statistics of the true prior distribution, *p*(***z***). For simplicity, we define the correct option in a perceptual task as the option associated with the largest hidden variable, *i*_correct_ = argmax_*i*_ *z*_*i*_, which, for example, can be interpreted as the highest contrast in a contrast discrimination task.

In both value-based and perceptual tasks, we assume that the decision-maker tries to maximize the expected reward under time constraints. Specifically, we focus on reaction time tasks in which the decision-maker is free to choose at any time within each trial, and proceeds through a long sequence of trials within a fixed time period. The total number of trials, and thus the total reward throughout the entire trial sequence, depends on how rapidly the decision-maker chooses in each trial: faster decisions allow for more of them in the same amount of time. However, due to noisy evidence, collecting more such evidence in each trial yields better choices, which results in a tradeoff between speed and accuracy.

### Optimal decision policy

We assume that the decision-maker’s aim is to maximize the total expected reward obtained in this task. The optimal decision policy comprises two key components: optimal online inference of the hidden variables by accumulating the evidence about them, and optimal rules for stopping the evidence accumulation to make a choice.

#### Optimal evidence accumulation

Here we provide a general formulation that includes correlations among options in the generative models. After some time *t* = *nδt*, the decision-maker’s posterior belief about the true hidden-variables *p*(***z***|*δ**x***_1_,…, *δ**x***_*n*_) is found by Bayes’ rule, 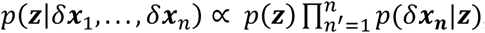, using the fact that *δ**x***_*n*′_ (*n*^′^ = 1,…, *n*) is independent and identically distributed (i.i.d.) across time. This results in

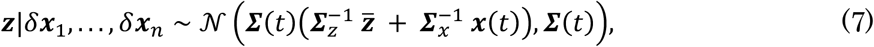

where we have defined 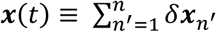 as the sum of all momentary evidence up to time *t*, and 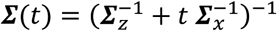 as the posterior covariance. The temporally accumulated evidence ***x***(*t*) and the time *t* provide the sufficient statistics for ***z***, and thus for the rewards ***r*** ≡ (*r*_1_,…, *r*_*N*_)^⊤^ associated with individual options. For the value-based case, the reward ***r*** equals the true hidden variable ***z***, that is ***r*** = ***z***, such that the expected option reward, 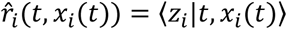, is the mean of the above posterior. For the perceptual case, the rewards associated with individual options are expressed as a vector ***r*** such that *r*_*i*_ = *r*_correct_ when *i* is the correct option and *r*_*i*_ = *r*_incorrect_ otherwise. Thus the expected reward for option *i* is *r*_*i*_(*t*, ***x***(*t*)) = *r*_correct_ *p*(*i* = *i*_correct_ | *t*, ***x***(*t*)) + *r*_incorrect_ *p*(*i* ≠ *i*_correct_ | *t*, ***x***(*t*)). Because *δ**x***_*n*_^′^ is i.i.d. in time, ***x***(*t*) is a random walk in an *N*-dimensional space (the thick black trace in Figure 2a). The next question is when to stop accumulating evidence and which option to choose at that point.

#### Optimal stopping rules

To find the optimal policy, we utilize tools from dynamic programming^17,18^. One such tool is the “value function” *V*(⋅), which can be defined recursively through Bellman’s equation. This value function returns for each state of the accumulation process (identified by the sufficient statistics) the total reward (including accumulation cost) the decision maker expects to receive from this state onward when following the optimal policy.

Let us first consider this value function for the case of a single choice, in which her aim is to maximize the expected reward for this choice minus some cost *c* per unit time for accumulating evidence (if there were no such cost, no decisions would ever be made). At any point in time *t*, the decision maker can either decide to make a choice, yielding the highest of the *N* expected rewards, or to accumulate more evidence for some small time *δt*, resulting in cost – *cδt*, and expected future reward given by the value function at time *t* + *δt*. By Bellman’s principle of optimality, the best action corresponds to the one yielding the highest expected reward, resulting in Bellman’s equation^17,18^

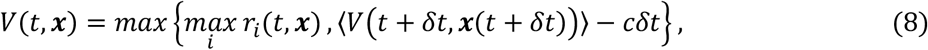

where the expected rewards *r*_*i*_(*t*, ***x***) differ between perceptual and value-based choices (see previous section; in both cases, they are functions of ***x*** and *t*), and the expectation in the second term is across expected changes of the accumulated evidence, *p*(***x***(*t* + *δt*) | ***x***(*t*), *t*). The intersection between the two terms within {⋅,⋅} determines the decision boundaries for stopping the evidence accumulation, and thus the optimal policy.

In more realistic setups, decision makers make a sequence of choices within a limited time period, in which case the aim of maximizing the total reward becomes equivalent (assuming long time periods) to maximizing their reward rate *ρ*, which is the expected reward for either choice divided by the expected time between consecutive choices. This reward rate is thus given by 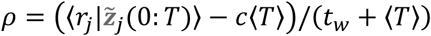, where *T* is the evidence accumulation time, *t*_*w*_ is the waiting time after choices (including the possible delays in motor responses) before onset of evidence for the next choice, and the expectation 〈⋅〉 here is across choices *j*. The value function associated with the reward-rate maximizing policy differs from the above by introducing an additional opportunity cost *ρ* per unit time. For immediate choices, this introduces the cost – *ρt*_*w*_ that the decision maker has to wait until the next trial (assuming 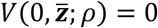, see below). For accumulating more evidence, the associated cost increases from – *cδt* to – (*c* + *ρ*)*δt*. Overall, this leads to Bellman’s equation Eq. (1) as given in the main text. If we set *ρ* = 0, we recover Bellman’s equation for single, isolated choices.

To find the optimal policy for the above cases numerically, we computed the value function by backward induction^20^, using Bellman’s equation. Bellman’s equation expresses the value function at time *t* as a function of the value function at time *t* + *δt*. Therefore, if we know the value function at some time *T*, we can compute it at time *T* − *δt*, then *T* − 2*δt*, and so on, until time *t* = 0. To find the reward rate, which is required to compute the value function, we initially set it to *ρ* = 0, computed the full value function, and then updated it iteratively by root finding until 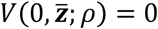, re-computing the full value function in each root finding step (see Drugowitsch et al., 2011^55^ for the rationale behind this procedure).

Unless otherwise mentioned, we used *T* = 10*s* and *δt* = 0.005*s* for all simulations. That is, we assumed *V*(*T* = 10, ***x***; *ρ*) to be given by the value for immediate choices, and then moved backwards in time in steps of 0.005s to find the value function by backward induction until *t* = 0. Furthermore, we set the prior parameters of the true, latent variables ***z*** to 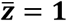 and ***∑***_***z***_ = ***I***. The waiting time was fixed to *t*_*w*_ = 0.5 s, and the accumulation cost to *c* = 0 (i.e., the opportunity cost *ρ* was the only cost). The results did not change qualitatively when changing the values of these parameters.

### Boundary structure analysis

Interestingly, we found that the decision boundaries in value-based tasks generally have a remarkable symmetry that reduces the optimal policy to a simple neural computation. All the decision boundaries are parallel to the diagonal — the line connecting (0,0,…, 0) and (1,1,…, 1).

In value-based tasks, this symmetry emerges from the fact that the state transition probability, *p*(***x***(*t*)|***x***(*t* + *δt*)), is invariant to translational shifts in ***x***. We can prove that the value function increases linearly along the diagonal, *V*(*t*, ***x*** + **1***C*) = *V*(*t*, ***x***) + *C* and **∇**_*x*_ *V*(*t*, ***x***) ≥ 0, where *C* is an arbitrary scalar. From these properties of the value function, we can prove that the decision boundaries are “parallel” to the diagonal: ∀*i*, *B*(*t*, *x*_*i*_ + *C*) = *B*(*t*, *x*_*i*_) + **1***C*, where *B*(*t*, *x*_*i*_) is a set of points that define for a fixed *x*_*i*_ the boundary in *x*_*j*≠*i*_ at which a decision ought to be made. The formal proofs are provided in **Supplementary Note 1**.

We can demonstrate the same symmetry in the perceptual tasks, even though it arises from a different mechanism: in perceptual tasks, by construction, the value function is determined by the probability of each option being the correct answer. Because this probability is already normalized such that the sum of all the probabilities across options is 1, the resulting value function is constant along the diagonal (in contrast to the value-based case in which the value function increases linearly along the diagonal). This yields the symmetry of decision boundaries along the diagonal.

### Circuit implementation of the optimal policy

It may seem difficult for biological systems to implement the optimal decision boundaries as these boundaries are, in general, represented by *N* time-dependent nonlinear functions *F*_*i*_(*t*, ***x***(*t*)) = 0 corresponding to the individual options, *i* = 1,…, *N*, that depends on *N* and other task contingencies. Fortunately, however, because of the symmetry of these boundaries (see main text), the decision policy effectively reduces to a lower dimensional representation (*N* − 1 dimensions for an *N*-alternative choice), which supports a simpler implementation of these boundaries. The key idea is as follows. The original decision policy representation assumes evidence accumulation by a simple random walk (diffusion) process in a linear space, which is terminated by a set of complex decision boundaries as a stopping rule. However, if we nonlinearly constrain the evidence accumulation space, we can vastly simplify these boundaries and instead can use constant decision thresholds that are independent across options.

More specifically, there exists a variable transformation, *ϕ*_*t*_: ***x***(*t*) ↦ ***x***^∗^(*t*) ≡ ***x*** + Δ_*x*_ ***1*** with a scalar Δ_*x*_, under which the optimal policy becomes equivalent to comparing each element 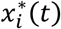 to a constant threshold *θ*_*x*_ satisfying 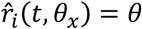. This variable transformation projects the states ***x*** onto an *N* − 1 dimensional manifold *M*_*θ*_ that is differentiable everywhere and asymptotically approaches the plane 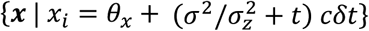 in the limit of ∀*j* ≠ *i*: *x*_*j*_ → −∞ for each *i*, where *σ*^2^ and 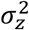 are the variances of likelihood and prior, respectively. The intersection of *M*_*θ*_ and the constant thresholds *x*_*i*_ = *θ*_*x*_ (∀*i*) implements effectively the same decision policy as the original one (see **Supplementary Note 2**).

Moreover, for some fixed time *t*, this manifold *M*_*θ*_ is well-approximated by the parameterized surface 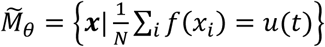, where *f*(*x*) is an arbitrary increasing, differentiable function that asymptotically approaches zero in the limit of *x*_*i*_ → −∞; *u*(*t*) is a scalar parameter. The variable transformation 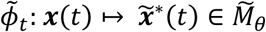 is achieved by a recurrent neural process shown in Figure 2c, which implements the following update rule,

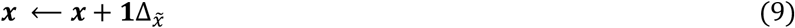

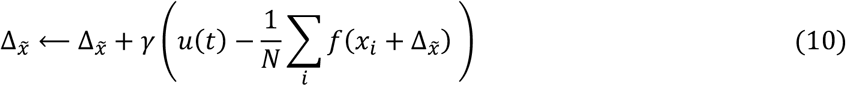

where *γ* is the update rate. Here, the second equation finds the appropriate 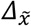, whereas the first equation performs the projection. This circuit comprises a nonlinear normalization of neural activities, 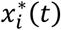, controlled by an “urgency signal”, *u*(*t*). Further, the circuit performs divisive normalization at a slower time-scale (see Equation 3, main text).

For subsequent simulations, we use the following sequence of discretized steps for each time-step of incoming momentary evidence: accumulate evidence according to Equation 2, project the newly accumulated evidence onto a nonlinear manifold by iterating Equation 4 (or 9-10) five times, perform divisive normalization as in Equation 3, add independent noise ξ_*i*_ on the individual output units or, equivalently, to the decision bounds (only for simulations corresponding to Figures 5-6). We follow this sequence because we assume that the projection happens at a much faster time-scale than divisive normalization (see main text). However, as we show in **Supplementary Note 5** and **Supplementary Figure S3**, this particular order of the time-discretized steps is inconsequential.

We found that a linear urgency signal, *u*(*t*) = *β t* + *u*_0_, approximates well the collapse of the optimal decision boundaries. Here, *β* and *u*_0_ are the slope and offset of the function, respectively, which we optimized in the subsequent simulations to maximize the reward rates. For the nonlinear function *f*, we used a rectified power function *f*(*x*_*i*_) = ⌊*x*_*i*_⌋^*α*^, with the exponent fixed to *α* = 1.5. The update rate of the projection in Equations 4, 10 was fixed to *γ* = 0.4. We also fixed the gain the divisive normalization term, *K*, to the mean reward across all trials and options, whereas *σ*_*h*_ was optimized. We ran the simulation for *T* = 10*s* with time-steps of *δt* = 0.005*s*. We identified the optimal parameters (i.e., the parameters that maximize the reward rate) with an exhaustive search for followed by a simplex optimization^56^. For *N* = 3 and *N* = 4, the circuit was confirmed to yield near-optimal reward rates for a reasonably wide range of the mean reward (from 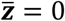 to 5).

### Violation of the IIA, similarity effect, and violation of the regularity principle

To simulate the third option effect violating IIA and regularity principles, and reproducing the similarity effect, we perform simulations to reoptimize our optimal neural circuit for *N* choice options with independent variability added to each accumulator at every time-step. We simulate the model for a fixed duration of *T* = 200*ms* as in Louie, et. al. (2013) ^15^ with time-steps of *δt* = 1*ms* and pick the option with the highest accumulator value at the end of the trial. The rewards for the three options were chosen uniformly from z_1_ ∊ [25,35], z_2_ = 30, z_3_ ∊ [0,30]. The momentary evidence was uncorrelated for IIA and regularity principles with ***∑***_*x*_ = *σ* 𝟙, whereas for the similarity effect, the momentary evidence for two of the choice options was positively correlated with correlation coefficient 0.1.

## References

1. Gold, J. I. & Shadlen, M. N. The Neural Basis of Decision Making. Annu. Rev. Neurosci. 30, 535–574 (2007).

2. Platt, M. & Glimcher, P. Neural correlates of decision variables in parietal cortex. Nature 400, 233–238 (1999).

3. Wang, X. J. Decision Making in Recurrent Neuronal Circuits. Neuron 60, 215–234 (2008).

4. Churchland, A. K. & Ditterich, J. New advances in understanding decisions among multiple alternatives. Curr. Opin. Neurobiol. 22, 920–926 (2012).

5. Ditterich, J. A comparison between mechanisms of multi-alternative perceptual decision making: Ability to explain human behavior, predictions for neurophysiology, and relationship with decision theory. Front. Neurosci. 4, 1–24 (2010).

6. Krajbich, I. & Rangel, A. Multialternative drift-diffusion model predicts the relationship between visual fixations and choice in value-based decisions. Proc. Natl. Acad. Sci. U. S. A. 108, 13852–13857 (2011).

7. Tajima, S., Drugowitsch, J. & Pouget, A. Optimal policy for value-based decision-making. Nat. Commun. 7, 12400 (2016).

8. Louie, K., Grattan, L. E. & Glimcher, P. W. Reward value-based gain control: divisive normalization in parietal cortex. J. Neurosci. 31, 10627–10639 (2011).

9. Louie, K., LoFaro, T., Webb, R. & Glimcher, P. W. Dynamic Divisive Normalization Predicts Time-Varying Value Coding in Decision-Related Circuits. J. Neurosci. 34, 16046–16057 (2014).

10. Churchland, A. K., Kiani, R. & Shadlen, M. N. Decision-making with multiple alternatives. Nat. Neurosci. 11, 693–702 (2008).

11. Louie, K., Khaw, M. W. & Glimcher, P. W. Normalization is a general neural mechanism for context-dependent decision making. Proc. Natl. Acad. Sci. U. S. A. 110, 6139–44 (2013).

12. Roe, R. M., Busemeyer, J. R. & Townsend, J. T. Multialternative decision field theory: a dynamic connectionist model of decision making. Psychol. Rev. 108, 370–392 (2001).

13. Furman, M. & Wang, X. J. Similarity Effect and Optimal Control of Multiple-Choice Decision Making. Neuron 60, 1153–1168 (2008).

14. Albantakis, L. & Deco, G. The encoding of alternatives in multiple-choice decision making. Proc. Natl. Acad. Sci. U. S. A. 106, 10308–10313 (2009).

15. Teodorescu, A. R. & Usher, M. Disentangling decision models: from independence to competition. Psychol. Rev. 120, 1–38 (2013).

16. Shadlen, M. N. & Shohamy, D. Decision Making and Sequential Sampling from Memory. Neuron 90, 927–939 (2016).

17. Drugowitsch, J., Moreno-Bote, R., Churchland, A. K., Shadlen, M. N. & Pouget, A. The cost of accumulating evidence in perceptual decision making. J. Neurosci. 32, 3612–28 (2012).

18. Mahadevan, S. Average reward reinforcement learning: Foundations, algorithms, and empirical results. Mach. Learn. 22, 159–195 (1996).

19. Bellman, R. E. Dynamic Programming. (1957).

20. Brockwell, A. E. & Kadane, J. B. A Gridding Method for Bayesian Sequential Decision Problems. J. Comput. Graph. Stat. 12, 566–584 (2003).

21. Baum, C. W. & Veeravalli, V. V. Sequential procedure for multihypothesis testing. IEEE Trans. Inf. Theory 40, 1994–2007 (1994).

22. Dragalin, V. P., Tartakovsky, A. G. & Veeravalli, V. V. Multihypothesis sequential probability ratio tests - Part II: accurate asymptotic expansions for the expected sample size. IEEE Trans. Inf. Theory 46, 1366–1383 (2000).

23. Bogacz, R. & Gurney, K. The basal ganglia and cortex implement optimal decision making between alternative actions. Neural Comput. 19, 442–477 (2007).

24. Carpenter, R. & Williams, M. Neural computation of log likelihood in control of saccadic eye movement. Nature 377, 59–62 (1995).

25. Brown, S. & Heathcote, A. A Ballistic Model of Choice Response Time. Psychol. Rev. 112, 117–128 (2005).

26. Thura, D. & Cisek, P. Deliberation and commitment in the premotor and primary motor cortex during dynamic decision making. Neuron 81, 1401–1416 (2014).

27. Thura, D. & Cisek, P. Modulation of Premotor and Primary Motor Cortical Activity during Volitional Adjustments of Speed-Accuracy Trade-Offs. J. Neurosci. 36, 938–956 (2016).

28. Keller, E. L. & McPeek, R. M. Neural discharge in the superior colliculus during target search paradigms. Ann. N. Y. Acad. Sci. 956, 130–42 (2002).

29. Hick, W. E. On the rate of gain of information in children. Q. J. Exp. Psychol. 4, 11–26 (1952).

30. Hyman, R. Stimulus information as a determinant of reaction time. J. Exp. Psychol. 45, 188–196 (1953).

31. Pastor-Bernier, A. & Cisek, P. Neural Correlates of Biased Competition in Premotor Cortex. J. Neurosci. 31, 7083–7088 (2011).

32. Mendonca, A. G. et al. The impact of learning on perceptual decisions and its implication for speed-accuracy tradeoffs. bioRxiv 501858 (2018). doi:10.1101/501858

33. Luce, R. D. Individual Choice Behavior: A Theoretical Analysis. (Wiley, New York, 1959).

34. Samuelson, P. A. Foundations of Economic Analysis. (Harvard University Press, Cambridge, MA, 1947).

35. Stephens, D. W. Foraging Theory. (Princeton University Press, Princeton, NJ, 1986).

36. Bateson, M., Healy, S. D. & Hurly, T. A. Context-dependent foraging decisions in rufous hummingbirds. Proc. Biol. Sci. 270, 1271–6 (2003).

37. Shafir, S., Waite, T. A. & Smith, B. H. Context-dependent violations of rational choice in honeybees (Apis mellifera) and gray jays (Perisoreus canadensis). Behav. Ecol. Sociobiol. 51, 180–187 (2002).

38. Tversky, A. & Simonson, I. Context-dependent Preferences. Manage. Sci. 39, 1179–1189 (1993).

39. Huber, J. et al. Adding Asymmetrically Dominated Alternatives: Violations of Regularity and the Similarity Hypothesis. J. Consum. Res. 9, 90–98 (1982).

40. Tversky, A. Elimination by aspects –A theory of choice. Psychol. Rev. 79, 281–299 (1972).

41. Gluth, S., Spektor, M. S. & Rieskamp, J. Value-based attentional capture affects multi-alternative decision making. Elife 7, (2018).

42. Tsetsos, K., Chater, N. & Usher, M. Salience driven value integration explains decision biases and preference reversal. Proc. Natl. Acad. Sci. U. S. A. 109, 9659–64 (2012).

43. Tsetsos, K., Moran, R., Moreland, J., Chater, N. & Usher, M. Economic irrationality is optimal during noisy decision making. Proc. Natl. Acad. Sci. U. S. A. 113, (2016).

44. Pettibone, J. C. Testing the effect of time pressure on asymmetric dominance and compromise decoys in choice. Judgm. Decis. Mak. 7, 513–521 (2012).

45. Trueblood, J. S., Brown, S. D. & Heathcote, A. The multiattribute linear ballistic accumulator model of context effects in multialternative choice. Psychol. Rev. 121, 179–205 (2014).

46. Usher, M. & McClelland, J. L. The time course of perceptual choice: The leaky, competing accumulator model. Psychol. Rev. 108, 550–592 (2001).

47. McMillen, T. & Holmes, P. The dynamics of choice among multiple alternatives. J. Math. Psychol. 50, 30–57 (2006).

48. Kveraga, K., Boucher, L. & Hughes, H. C. Saccades operate in violation of Hick’s law. Exp. Brain Res. 146, 307–314 (2002).

49. Lawrence, B. M., St John, A., Abrams, R. a & Snyder, L. H. An anti-Hick’s effect in monkey and human saccade reaction times. J. Vis. 8, 26.1–7 (2008).

50. Treisman, A. & Souther, J. Search asymmetry: a diagnostic for preattentive processing of separable features. J. Exp. Psychol. Gen. 114, 285–310 (1985).

51. Steverson, K., Brandenburger, A. & Glimcher, P. W. Rational Imprecision: Information-Processing, Neural, and Choice-Rule Perspectives. SSRN Electron. J. (2017). doi:10.2139/ssrn.3056332

52. Bogacz, R., Usher, M., Zhang, J. & McClelland, J. L. Extending a biologically inspired model of choice: multi-alternatives, nonlinearity and value-based multidimensional choice. Philos. Trans. R. Soc. Lond. B. Biol. Sci. 362, 1655–1670 (2007).

53. Simonson, I. Choice based on reasons: The case of attraction and compromise effects. J. Consum. Res. 16, 158–174 (1989).

54. Howes, A., Warren, P. A., Farmer, G., El-Deredy, W. & Lewis, R. L. Why contextual preference reversals maximize expected value. Psychol. Rev. 123, 368–391 (2016).

55. Drugowitsch, J., Moreno-Bote, R. & Pouget, A. Optimal decision bounds for probabilistic population codes and time varying evidence. Nat. Preced. (2011). doi:10.1038/NPRE.2011.5821.1

56. Acerbi, L. & Ji, W. Practical Bayesian Optimization for Model Fitting with Bayesian Adaptive Direct Search. 1836–1846 (2017).

