## Supplementary Material for "Optimal policy for multi-alternative decisions"

Supplementary Information

### Supplementary Notes

#### 1 Structure of the value function and the optimal decision boundaries

##### *The value function*

In this section, we provide an analytic characterization of the decision boundary structure. To do so, we focus on the value function in the single-choice value-based decision tasks; the result for the reward rate case is not shown, but follows a similar analysis. Assume that $\boldsymbol{X}(t)$ is the stochastic process (or "decision variable") that describes the expected reward in $N$-dimensional space. Furthermore, assume that $\boldsymbol{X}(t)$ is shift-invariant, that is $\boldsymbol{X}(\tau) \mid (\boldsymbol{X}(t) + \boldsymbol{C}) = \left(\boldsymbol{X}(\tau) \mid \boldsymbol{X}(t)\right) + \boldsymbol{C}$, where $\tau \geq t$. For simple (even correlated) setups, this will hold. In particular, it holds for all cases discussed in the main text.

In this context, the value function is non-recursively given by

$$V(t, \boldsymbol{x}) = \max_{\tau \geq t} \left\langle \max_i X_i(\tau) - c(\tau - t) \middle| \boldsymbol{X}(t) = \boldsymbol{x} \right\rangle, \tag{1}$$

where the expectation is over the time-evolution of $\boldsymbol{X}$.

Below we show the value function to have the following properties:

1. $V(t, \boldsymbol{x} + \boldsymbol{1}\, C) = V(t, \boldsymbol{x}) + C$.
2. $V(t, \boldsymbol{x})$ is increasing in each element of $\boldsymbol{x}$.
3. $V(t, \boldsymbol{x}) \leq V(t, \boldsymbol{x} + \boldsymbol{e}_i\, C) \leq V(t, \boldsymbol{x}) + C$, where $\boldsymbol{e}_i$ is the $i$th basis vector of a Cartesian basis.
4. $V(t, \boldsymbol{x}) + \min_i C_i \leq V(t, \boldsymbol{x} + \boldsymbol{C}) \leq V(t, \boldsymbol{x}) + \max_i C_i$, where $C_i$ is the $i$ th element of $\boldsymbol{C}$.

Property 2 implies that $V(t, \boldsymbol{x})$ is continuous and differentiable. Thus, this property can be expressed as $\boldsymbol{\nabla}_x V(t, \boldsymbol{x}) \geq 0$, where the inequality is on each element of the gradient separately. As $C$ in property 3 can be arbitrarily small, it is a generalization of property 2, such that we only need to show property 3. Property 1 is a special case of property 4 in which $\boldsymbol{C} = \boldsymbol{1}\, C$, such that $\min_i C_i = \max_i C_i = C$.

##### Property 1

Fix some stopping times $\tau_1, \ldots, \tau_N$. Then, the value function at time $t$ is given by

$$\left\langle \sum_i 1_{\tau_i < \mathrm{mi}\ _{j \neq i} \tau_j} X_i(\tau_i) - c(\min_i \tau_i - t) \middle| \boldsymbol{X}(t) = \boldsymbol{x} \right\rangle, \tag{2}$$

where the indicator function $1_a$ is 1 if $a$ is true, and 0 otherwise. Thus, if we set the starting point to $\boldsymbol{x} + \boldsymbol{1}\, C$, we find

$$\left\langle \sum_i 1_{\tau_i < \min_{j\neq i} \tau_j} X_i(\tau_i) - c\left(\min_i \tau_i - t\right) \middle| \boldsymbol{X}(t) = \boldsymbol{x} + \boldsymbol{1}C \right\rangle$$

$$= \left\langle \sum_i 1_{\tau_i < \min_{j\neq i} \tau_j} (X_i(\tau_i) + C) - c\left(\min_i \tau_i - t\right) \middle| \boldsymbol{X}(t) = \boldsymbol{x} \right\rangle$$

$$= \left\langle \sum_i 1_{\tau_i < \min_{j\neq i} \tau_j} X_i(\tau_i) - c\left(\min_i \tau_i - t\right) \middle| \boldsymbol{X}(t) = \boldsymbol{x} \right\rangle + C, \quad (3)$$

where the last line follows because the indicator function is only 1 for a single $n$. This is true for all choices of stopping times, and so also for the maximum over stopping times and choices.

##### Properties 2 and 3

Fix some integer $k$ and stopping times $\tau_1, \dots, \tau_N$. For starting point $\boldsymbol{x} + \boldsymbol{e}_k C$ we get

$$\left\langle \sum_i 1_{\tau_i < \min_{j\neq i} \tau_j} X_i(\tau_i) - c\left(\min_i \tau_i - t\right) \middle| \boldsymbol{X}(t) = \boldsymbol{x} + \boldsymbol{e}_k C \right\rangle$$

$$= \left\langle \sum_{i\neq k} 1_{\tau_i < \min_{j\neq i} \tau_j} \left(X_i(\tau_i)\right) + 1_{\tau_k < \min_{j\neq k} \tau_j} (X_k(\tau_k) + C) - c\left(\min_i \tau_i - t\right) \middle| \boldsymbol{X}(t) = \boldsymbol{x} \right\rangle$$

$$= \left\langle \sum_i 1_{\tau_i < \min_{j\neq i} \tau_j} X_i(\tau_i) - c\left(\min_i \tau_i - t\right) \middle| \boldsymbol{X}(t) = \boldsymbol{x} \right\rangle + 1_{\tau_k < \min_{j\neq k} \tau_j} C, \quad (4)$$

Note that, for the last term of the last line, $0 \leq 1_{\tau_k < \min_{j\neq k} \tau_j} C \leq C$, which upper-bounds the increase by $C$. The above again holds for an arbitrary set of stopping times, such that it also holds for the maximum over stopping times and choices.

##### Property 4

Following the same argument as in the preceding sections, we find for initial state $\boldsymbol{x} + \boldsymbol{C}$ and fixed stopping times that the value function is given by

$$\left\langle \sum_i 1_{\tau_i < \min_{j\neq i} \tau_j} X_i(\tau_i) - c\left(\min_i \tau_i - t\right) \middle| \boldsymbol{X}(t) = \boldsymbol{x} \right\rangle + \sum_i 1_{\tau_i < \min_{j\neq i} \tau_j} C_i, \quad (5)$$

The last term is bounded by $\min_i C_i \leq \sum_i 1_{\tau_i < min_{j\neq i} \tau_j} C_i \leq \max_i C_i$, such that the result follows.

#### *Characterizing the optimal decision boundaries*

In this section we derive a few properties of the optimal decision boundaries, based on the above value function properties.

#### The expression for the optimal decision boundaries

Note that $V(t,\boldsymbol{x}) \geq \max_i x_i$. Furthermore, the decision maker ought to accumulate more evidence as long as $V(t,\boldsymbol{x}) > \max_i x_i$ and decide as soon as $V(t,\boldsymbol{x}) = \max_i x_i$. Let us assume that $x_1 > \max_{j>1} x_j$, such that, in case of a choice, option 1 ought to be chosen. The argument that follows is valid for all options, but we focus on option 1 for notational convenience. In this case, we have $x_j < x_1$ for all $j > 1$, and $V(t,\boldsymbol{x}) \geq x_1$. Furthermore, we accumulate evidence as long as $V(t,\boldsymbol{x}) > x_1$, and choose option 1 as soon as $V(t,\boldsymbol{x}) = x_1$. Note that $V(t,\boldsymbol{x})$ is increasing in $x_{2:N} \equiv x_2, \dots, x_N$, such that we will have $V(t,\boldsymbol{x}) > x_1$ for large $x_{2:N}$. Lowering $x_{2:N}$ will cause $V(t,\boldsymbol{x})$ to reduce until it reaches its lower bound, $V(t,\boldsymbol{x}) = x_1$, which is the point at which a decision ought to be made. Thus, the decision boundary is the "largest" $x_{2:N}$ (assuming natural vector ordering) at which $V(t,\boldsymbol{x}) = x_1$, or

$$B_1(t,x_1) \equiv \max \{x_{2:N} < x_1 \mid V(t,\boldsymbol{x}) = x_1\}, \tag{6}$$

where $x_{2:N} < x_1$ here denotes $x_j < x_1$ for all $j > 1$. This $B_1(t,x_1)$ is a set of points that, for a fixed $x_1$, define the boundary in $x_{2:N}$ at which a decision ought to be made. The above argument and resulting expression is valid for the decision boundaries associated with all options.

#### The decision boundaries are continuous and decreasing

To show that the decision boundaries are continuous, fix again $x_1$ such that $x_1 > \max_{j>1} x_j$. Furthermore, pick some $x_2$ and $x_2 + \delta$ that are both part of the vector elements of $B_1(t,x_1)$ (this restriction is necessary, as we cannot arbitrarily increase $x_2$ and still guarantee it to be part of the decision boundary). As the decision boundary is determined by the largest $x_{2:N}$ such that $V(t,\boldsymbol{x}) = x_1$, increasing $x_2$ while leaving all other elements constant will cause $V(t,\boldsymbol{x} + \boldsymbol{e}_2\,\delta) > x_1$. Therefore, we need to reduce another element of $x_{2:N}$ such that $V(t,\boldsymbol{x}) = x_1$ is again satisfied. As $\delta$ is arbitrarily small and $V(t,\boldsymbol{x})$ is increasing in all elements of $\boldsymbol{x}$, the decision boundary is continuous. Furthermore, as increasing one element of $x_{2:N}$ causes a decrease in other elements, the decision boundary as function of $x_2$ is decreasing in $x_{3:N}$.

#### The decision boundaries are "parallel" to the diagonal

Let us add a constant vector $\mathbf{1}C$ to all elements in $B_1(t,x_1)$. Defining $\boldsymbol{x}' = \boldsymbol{x} + \mathbf{1}C$, this results in

$$\begin{aligned} B_1(t,x_1) + \mathbf{1}C &= \max \{x_{2:N} < x_1 \mid V(t,\boldsymbol{x}) = x_1\} + \mathbf{1}C \\ &= \max \{x'_{2:N} < x'_1 \mid V(t,\boldsymbol{x}' - \mathbf{1}C) = x'_1 - C\} \\ &= \max \{x'_{2:N} < x'_1 \mid V(t,\boldsymbol{x}') = x'_1\} \\ &= B(t,x'_1) \\ &= B(t,x_1 + C). \end{aligned} \tag{7}$$

Thus, $B_1(t,\mathrm{x}_1 + C) = B_1(t,\mathrm{x}_1) + \mathbf{1}C$ which implies that the decision boundaries are parallel to the diagonal.

This implies that, for decision-making, only the accumulation space orthogonal to the direction given by $\mathbf{1}$ matters. Mapping onto this space could be achieved by $y_i = x_i - (N-1)^{-1}\sum_{j\neq i} x_j$ , or other arbitrary projections on $N-1$ dimensional manifolds, which maps the accumulation into an $N-1$ dimensional subspace.

#### 2 Neural circuit implementation of the decision policy

In this section, we describe step-by-step the reason why the proposed recurrent neural circuit can approximate the optimal decision policy for $N$-alternative value-based decisions. Again, here we focus on the single-choice value-based decision tasks; the same arguments hold for the reward rate cases.

##### *Decision boundaries as a set of manifold intersections*

The optimal decision boundaries are determined by Bellman's equation,

$$V(t, \boldsymbol{x}) = \max\left\{\max_i \hat{r}_i(t, x_i), \langle V(t + \delta t, \boldsymbol{x})\rangle - c\,\delta t\right\}, \tag{8}$$

In the curled bracket, the first term corresponds to the value for deciding and choosing something right now, whereas the second term corresponds to the value for waiting (postponing the decision) to accumulate more evidence. Let us fix some time $t$. For this fixed time, the boundaries $B$ between deciding and waiting are defined as a set of states where those two value functions equal to each other, i.e.,

$$B \equiv \left\{\boldsymbol{x} \,\middle|\, \max_i \hat{r}_i(t, x_i) = \langle V(t + \delta t, \boldsymbol{x})\rangle - c\,\delta t\right\}, \tag{9}$$

which is described as a set of intersections between the following two $N - 1$ dimensional manifolds,

$$L_\theta \equiv \left\{\boldsymbol{x} \,\middle|\, \max_i \hat{r}_i(t, x_i) = \theta\right\},$$

$$M_\theta \equiv \{\boldsymbol{x} \mid \langle V(t + \delta t, \boldsymbol{x})\rangle - c\,\delta t = \theta\}, \tag{10}$$

with a scalar reward parameter $\theta$ varied from $-\infty$ to $\infty$. $L_\theta$ and $M_\theta$ represent the level sets of value functions for choosing either option right now and for waiting to accumulating more evidence, respectively. Just as $B$, $L_\theta$ and $M_\theta$ are defined for some fixed time $t$. As shown in **Figure S1a** and **S1b**, $L_\theta$ represents one corner of an $N$-dimensional hypercube (i.e., an orthant, as described later) that is intersected by $M_\theta$. The point of this intersection corresponds to the part of the decision boundary that promises reward $\theta$. Therefore, the complete set of decision boundaries $B$ can be expressed as a "chain" of intersections between the two manifolds $L_\theta$ and $M_\theta$, ordered by the reward parameter $\theta$ (**Figure S1c**):

$$B = \{B_\theta \mid -\infty < \theta < \infty\} \tag{11}$$

$$B_\theta \equiv L_\theta \cap M_\theta. \tag{12}$$

For each $\theta$ the dimensionality of $B_\theta$ is $N - 2$ because it is an intersection of two $N - 1$ dimensional manifolds, which makes the full decision boundary $B$ an $N - 1$ dimensional manifold. Recall that all the value functions are shift-invariant in the dimension parallel to the diagonal, and that the set of decision boundaries, $B$, is "parallel" to the diagonal. Using this fact, $B$ can be expressed in a different way as follows:

$$B = \{B_{\theta+\Delta_\theta} \mid -\infty < \Delta_\theta < \infty\}$$

$$= \{B_\theta + \mathbf{1}\Delta_\theta \mid -\infty < \Delta_\theta < \infty\}, \quad (13)$$

where $+\mathbf{1}\Delta_\theta$ represents a translational shift of the set along the diagonal vector, $\mathbf{1}$, with a distance $\Delta_\theta$. Thus, rather than defining the set of all boundaries by the intersection $B_\theta$ between $L_\theta$ and $M_\theta$ for all reward levels $\theta$ (first line), we can define it as one such intersection $B_\theta$ for some arbitrary fixed $\theta$, translated in directions of the diagonal $\mathbf{1}$ (second line).

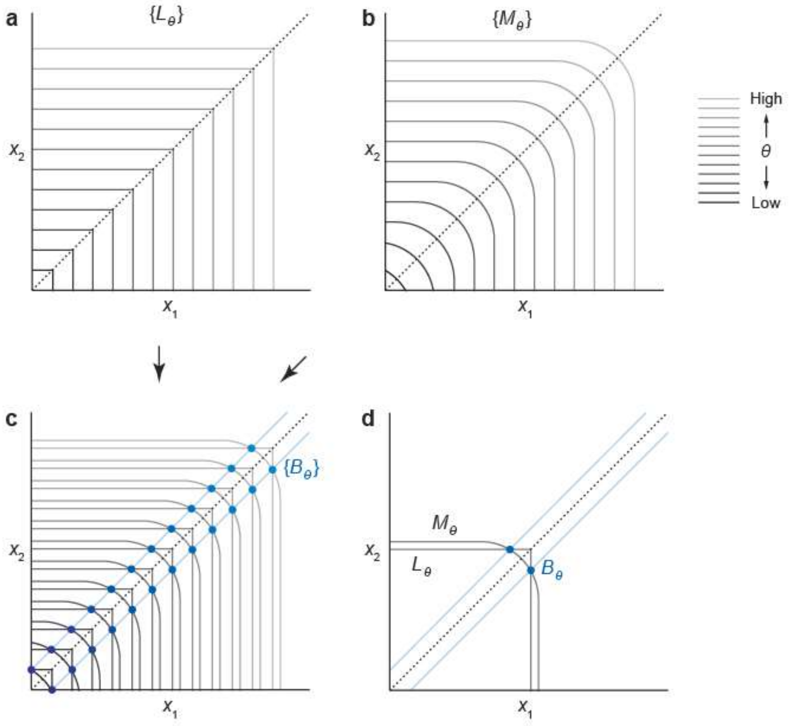


**Supplementary Figure S1**

Schematic illustrations of the decision boundaries defined by manifold intersections. A two-alternative case is shown for the visualization purpose although the same argument applies to arbitrary $N$-alternative problems. (a) The manifold set $\{L_\theta\}$, which describes the value function for "deciding right now." (b) The manifold set $\{M_\theta\}$, which describes the value function for "waiting to accumulate more evidence." (c) The set of decision boundaries, $B \equiv \{B_\theta | -\infty < \theta < \infty\}$, is defined as a set of intersections of $L_\theta$ and $M_\theta$. (d) Because the decision boundaries defined by different $\theta$ are all symmetric along the diagonal (the dashed line), we can consider a lower-dimensional projection by fixing $\theta$.

##### *Constrained states*

As a next step we demonstrate that, if we restrict our evidence accumulation process state $\boldsymbol{x}$ to its projection $\boldsymbol{x}^*$ parallel to the diagonal $\mathbf{1}$ on the manifold $M_\theta$, then we can make optimal choices as soon as this projected state reaches the manifold $L_\theta$, which implies reaching $B_\theta = L_\theta \cap M_\theta$ (see also **Figure S1d**). For now we assume some arbitrary fixed $\theta$, but will later discuss that the argument is valid for any $\theta$. More formally, fix some arbitrary $\theta$ (and some time $t$, as in the previous section) and consider the following map:

$$\phi_\theta : \mathbb{R}^N \to M_\theta, \tag{14}$$

$$\phi_\theta : \boldsymbol{x} \mapsto \boldsymbol{x}^* \equiv \boldsymbol{x} + \mathbf{1}\Delta_x. \tag{15}$$

which projects each state along the diagonal onto manifold $M_\theta$. We call $\boldsymbol{x}^*$ the "constrained state." For a particular state $\boldsymbol{x} \in M_{\theta+\Delta_\theta}$ the extent $\Delta_x$ of this projection corresponds to $\Delta_x = -\Delta_\theta$, which yields the set of states that project into $B_\theta$ to be given by

$$\begin{aligned} \{\boldsymbol{x} \mid \exists \Delta_\theta : \boldsymbol{x} \in B_\theta + \mathbf{1}\,\Delta_x\} &= \{\boldsymbol{x} \mid \boldsymbol{x}^* \in B_\theta\} \\ &= \{\boldsymbol{x} \mid \boldsymbol{x}^* \in L_\theta\} \\ &= \left\{\boldsymbol{x} \,\middle|\, \max_i \hat{r}_i\,(t, x_i^*) = \theta\right\}, \end{aligned} \tag{16}$$

where the first equality follows from the definition of the projection, the second from the definition of $B_\theta$ as the intersection of $M_\theta$ and $L_\theta$ (recall that $\boldsymbol{x}^*$ is in $M_\theta$ by definition), and the third from the definition of $L_\theta$. Note that by Equation (13) the states that project into $B_\theta$ form the set of all decision boundaries $B$, such that we can re-express the above as

$$\{\boldsymbol{x} \mid \boldsymbol{x} \in B\} = \left\{\boldsymbol{x} \,\middle|\, \max_i \hat{r}_i(t, x_i^*) = \theta\right\}, \tag{17}$$

showing that, as long as the accumulation process is constrained to states in $M_\theta$, the decision boundary is formed by points on $L_\theta$.

In the value-based case, as $\hat{r}_i(t, x_i)$ is an increasing function of $x_i$ for each $i$, there exists a unique scalar $\theta_x \in \mathbb{R}$ such that $\hat{r}_i(t, \theta_x) = \theta$, with which the decision boundaries are described as

$$\{\boldsymbol{x} \mid \boldsymbol{x} \in B\} = \left\{\boldsymbol{x} \,\middle|\, \max_i x_i^* = \theta_x\right\}. \tag{18}$$

This equation shows that evaluating whether the state $\boldsymbol{x}$ hits a decision boundary or not is equivalent to evaluating whether the largest component of the constrained state $\boldsymbol{x}^*$ equals $\theta_x$ or not. Since the choice of $\theta$ is arbitrary, we can choose any $\theta$ that makes $\theta_x$ constant over time. With such a time-invariant $\theta_x$, the decision policy is implemented simply by evaluating whether the largest component of $\boldsymbol{x}^*$ exceeds a fixed threshold. Note also that, because the value function is decreasing for each element $x_i$ as described previously (**Supplementary Note 1**), the manifold $M_\theta$ is also a decreasing function for each element, thus the projection of the states $\boldsymbol{x}$ to $M_\theta$ is generally described as a mutual inhibition among the elements $x_i$ corresponding to the individual options.

##### *The structure of $L_\theta$ and $M_\theta$*

As a next step, we investigate the structure of the two manifolds in order to find functional forms that capture the symmetry of those manifolds.

As already described further above $L_\theta$ is a set of $N-1$ dimensional half-planes, $\{\boldsymbol{x} | x_i = \theta_x, x_{j\neq i} \leq \theta_x\}$ $(i = 1, \ldots, N)$. These half-planes collectively form the sides of an $N$-dimensional orthant whose origin is at $\theta_x \mathbf{1} = (\theta_x, \theta_x, \ldots, \theta_x)$ (**Figure S1d**). Due to this straightforward form, $L_\theta$ does not need to be approximated.

On the other hand, $M_\theta$ is a surface of a "smoothed orthant," which we define here as a differentiable $N-1$ dimensional manifold that asymptotically approaches $L_{\theta'}$ $(\exists\theta')$ in the limit of $\forall j \neq i : x_j \to -\infty$ for each $i$ (**Figure S1d**); this is because when all the options except for option $i$ have infinitely low values, the decision-maker should choose option $i$, which makes the value for waiting equal the value for choosing option $i$ minus the cost of time. In particular, from the Bellman equation and the Bayes rule applied to our setup, $\theta'$ could be defined explicitly as

$$\theta' = \theta_x + \left(\frac{\sigma^2}{\sigma_z^2} + t\right) c\,\delta t, \tag{19}$$

where $\sigma^2$ and $\sigma_z^2$ are the variances of the evidence noise and the prior, respectively. Using the symmetry along the diagonal line,

$$L_{\theta'} = L_{\theta_x} + \left(\frac{\sigma^2}{\sigma_z^2} + t\right) c\,\delta t\,\mathbf{1}, \tag{20}$$

This equation implies that $L_{\theta'}$ and thus $M_\theta$ move along the diagonal as time elapses.

Due to the symmetry, $M_\theta$ is invariant to permutations of the coordinates, $1, \ldots, N$. Thus, at the point $\boldsymbol{x} \in M_\theta$ such that $\forall\, i, j : x_i = x_j$, $M_\theta$ is orthogonal to the $N$-dimensional diagonal line, $\{\boldsymbol{x} | \forall\, i, j : x_i = x_j\}$. Note that $M_\theta$ has only one intersection with the diagonal line due to the decreasing property as we have described further above. Furthermore, as understood intuitively, if option $j$'s value is very low, the problem becomes effectively a comparison among the remaining $N-1$ options, $\{1, \ldots, N\} \setminus$ j. Because we have the same symmetry as before but now among those $N-1$ options; i.e., in the limit of $\forall\, j : x_j \to -\infty$, $M_\theta$ is orthogonal to the vector $\mathbf{1}^{\setminus j}$ that is defined by $\mathbf{1}_i^{\setminus j} = 1 - \delta_{ij}$ with Kronecker's delta, when $\boldsymbol{x} \in M_\theta$ satisfies $\forall\, i, i' \neq j : x_i = x_{i'}$. Similarly, if two options $j$ and $j'$ have infinitely low values, the effective problem becomes to compare the remaining $N-2$ options, $\{1, \ldots, N\} \setminus \{j, j'\}$, then the manifold is orthogonal to $\mathbf{1}^{\setminus\{j,j'\}}$ (where $\mathbf{1}_i^{\setminus\{j,j'\}} = 1 - \delta_{ij}\delta_{ij'}$) when $\boldsymbol{x} \in M_\theta$ satisfies $\forall\, i, i' \notin \{j, j'\} : x_i = x_{i'}$. Repeating the same argument reveals the whole hierarchy of symmetries in the manifold $M_\theta$. Note that when $\forall\, j \neq i : x_j \to -\infty$, the problem reduces to choosing from only one option $i$; at this limit, $M_\theta$ is orthogonal to $\mathbf{1}^{\setminus(\{1,\ldots,N\}\setminus i)} = \boldsymbol{e}_i$, agreeing with the aforementioned property that $M_\theta$ asymptotically approaches $L_{\theta'}(\exists\theta')$.

These properties of $M_\theta$ are well-captured by a manifold defined as follows:

$$\widetilde{M}_\theta = \left\{ \boldsymbol{x} \,\middle|\, \tfrac{1}{N}\textstyle\sum_i f(x_i) = u \right\}, \tag{21}$$

where $f(x_i)$ is an arbitrary increasing, differentiable function that asymptotically approaches zero in the limit of $x_i \to -\infty$. Here, $u$ is a scalar value, which generally increases with elapsed time to capture the time-dependent property of $M_\theta$. Moreover, by varying the functional form of $f$ and the value of parameter $u$, we can make $\widetilde{M}_\theta$ have the same position and curvature as $M_\theta$ at its intersection with the diagonal line, $\{\boldsymbol{x} \mid \boldsymbol{x} \in \widetilde{M}_\theta,\ \forall i, j : x_i = x_j\}$. This indicates that $\widetilde{M}_\theta$ can be a good approximation of $M_\theta$ around its intersection with the diagonal line, and thus $\widetilde{B}_\theta \equiv L_\theta \cap \widetilde{M}_\theta$ approximates $B_\theta$ well at points close to the diagonal line. The approximation around the diagonal line is particularly important because the assumed unbiased prior over rewards requires the initially expected rewards (at $t = 0$, before accumulating any evidence) to be symmetric across options, such that the decision variable $\boldsymbol{x}(t)$ fluctuates around the diagonal line.

##### *A recurrent circuit that approximates the constraining manifold*

We design a neural mechanism that constrains the neural population activity that encodes evidence accumulation to the manifold $\widetilde{M}_\theta$. Consider a map that projects each state along the diagonal onto the manifold $\widetilde{M}_\theta$, as follows:

$$\tilde{\phi}_\theta : \mathbb{R}^N \to \widetilde{M}_\theta, \tag{22}$$

$$\tilde{\phi}_\theta : \boldsymbol{x} \mapsto \tilde{\boldsymbol{x}}^* \equiv \boldsymbol{x} + \mathbf{1}\Delta_{\tilde{x}}, \tag{23}$$

Based on the arguments in the previous sections (Supplementary Note 1), the decision boundary $\{\boldsymbol{x} \mid \boldsymbol{x} \in B\}$ is approximated by $\left\{\boldsymbol{x} \,\middle|\, \max_i \tilde{x}_i^* = \theta_x \right\}$. That is, evaluating whether the state x hits a decision boundary or not is equivalent to evaluating whether the largest component of the constrained state $\tilde{x}^*$ equals $\theta_x$ or not. Again, $\widetilde{M}_\theta$ also depends on time, thus so does the map $\tilde{\phi}_\theta$. As shown in the previous section, the map $\tilde{\phi}_\theta$ can be implemented by a circuit that computes $\Delta_{\tilde{x}}$ satisfying the following property:

$$\tilde{\boldsymbol{x}}^* \in \widetilde{M}_\theta \Leftrightarrow \tfrac{1}{N}\textstyle\sum_i f(\tilde{\boldsymbol{x}}_i) = u, \tag{24}$$

$$\Leftrightarrow u - \tfrac{1}{N}\textstyle\sum_i f(x_i + \Delta_{\tilde{x}}) = 0. \tag{25}$$

If we define $E \equiv \frac{1}{2}\left(u - \frac{1}{N}\sum_i f(x_i + \Delta_{\tilde{x}})\right)^2$ its gradient is given by

$$-\frac{\partial E}{\partial \Delta_{\tilde{x}}} = \left(\frac{1}{N}\sum_i f'(x_i + \Delta_{\tilde{x}})\right)\left(u - \frac{1}{N}\sum_i f(x_i + \Delta_{\tilde{x}})\right). \tag{26}$$

Note that the first term on the left hand side of the equation is always zero or positive because $f$ is an increasing function as described in the previous section. Therefore, the following update rule is able to find $\Delta_{\tilde{x}}$ by approximate gradient descent:

$$\Delta_{\tilde{x}} \leftarrow \Delta_{\tilde{x}} + \gamma\left(u - \frac{1}{N}\sum_i f(x_i + \Delta_{\tilde{x}})\right), \tag{27}$$

where $\gamma$ is a small positive scalar that determines the update rate. The update process terminates when the term in the parenthesis becomes zero. This update rule is implemented by the recurrent neural circuit with activity normalization (implementing the projection) and urgency signal (realizing the time-variant nature of $M_\theta$) as mentioned in the main text.

Corresponding to the update of $\Delta_{\tilde{x}}$, each neuron's output is updated as follows:

$$f(\tilde{x}_i) \leftarrow f(\tilde{x}_i + \Delta_{\tilde{x}}). \tag{28}$$

Given that $f$ is invertible in $\tilde{x}_i \geq 0$, this update is the same as $\tilde{x}_i \leftarrow \tilde{x}_i + \Delta_{\tilde{x}}$. In our implementation, the projection was performed by applying Eqs. (27) and (28) 5 times for each evidence input at time $t$, assuming that the relaxation of neural activity is faster compared to the time scale of the evidence dynamics. Within every time step $t$, $f(\tilde{x}_i)$ was after the projection compared to a constant threshold $\theta \equiv f(\theta_x)$, which, for the monotonically increasing $f$, is equivalent to comparing $\tilde{x}_i$ with $\theta_x$.

#### 3 Experimental predictions

Here we provide a list of detailed predictions derived from our theoretical results. All of them can be tested with neurophysiological or behavioral experiments. We consider human or animal subjects performing a standard $N$-alternative value-based decision-making task with a reaction-time paradigm, in which the subjects choose one of $N$ options at their own pace, while trying to maximize the total reward within a session of fixed duration (i.e., they can make more choices if each of them is faster).

Suppose that we record the activity of a decision-related neuronal population (serially or simultaneously) during the task. We denote the entire population state by $\boldsymbol{x}(t) = \big(x_1(t), \dots, x_D(t)\big)$, where $D$ is the number of recorded neurons.

##### *Physiological predictions*

Here we provide a list of detailed predictions derived from our theoretical results. All of them can be tested with neurophysiological or behavioral experiments. We consider human or animal subjects performing a standard $\mathrm{N}$-alternative value-based decision-making task with a reaction-time paradigm, in which the subject choose one of $\mathrm{N}$ options at their own pace, while trying to maximize the total reward within a session of fixed duration (i.e., they can make more choices if each of them is faster).

Suppose that we record the activity of a decision-related neuronal population (serially or simultaneously) during the task. We denote the entire population state by $x(t) = \big(x_1(t), \dots, x_{\mathrm{D}}(t)\big)$, where $\mathrm{D}$ is the number of recorded neurons.

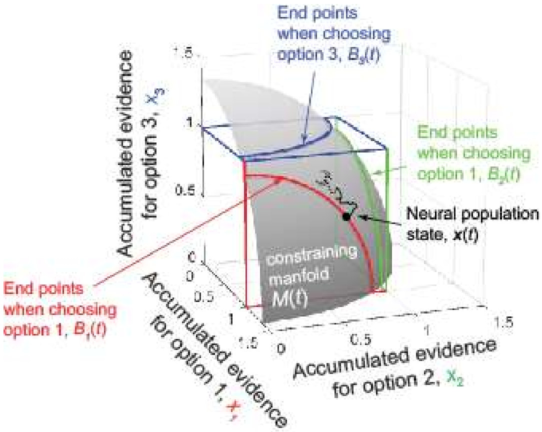


**Supplementary Figure S2. The manifold constraining neural population state.**

#### Predictions about the dynamics of evidence accumulation

1. The neural population activity is constrained on a low-dimensional manifold: at each time point $t$, the neural population state $\boldsymbol{x}(t)$ is constrained on an $N-1$ dimensional manifold, $\mathcal{M}(t)$ (the 'constraining manifold', the gray surface in **Supplementary Figure S2**).

2. The $N-1$ dimensional manifold is nonlinear: although the topological dimensionality (i.e., a locally defined dimensionality) of the constraining manifold $\mathcal{M}(t)$ is $N-1$, $\mathcal{M}(t)$ is curved and embedded within an $N$-dimensional space.

3. The $N-1$ dimensional manifold evolves over time: the constraining manifold $\mathcal{M}(t)$ varies over time, meaning that the $N-1$ dimensional manifold can only be observed for a fixed time. Otherwise, we would only observe an $N$-dimensional structure, resulting from the $N-1$ dimensional manifold being smeared out over time.

4. The effects of prior belief: the position and the shape of the constraining manifold $\mathcal{M}(t)$ depend on the prior knowledge about the value distribution (e.g., mean value over trials), but the dimensionality is always $N-1$.

5. The stability against the trial-to-trial option contingency: the constraining manifold $\mathcal{M}(t)$ is invariant to the option values within each trial. That is, if we compare a trial set with $N$ high-valued options against another trial set in which low and high values are mixed, the neural state trajectories can differ between those trial sets, but all the trajectories are constrained on the same $N-1$ dimensional manifold $\mathcal{M}(t)$.

6. The constraining manifold $\mathcal{M}(t)$ has a hierarchical symmetry as follows: on $\mathcal{M}(t)$, $x_i(t)$ is a decreasing function of $x_j(t)$ for all $i \neq j$. In particular, $\mathcal{M}(t)$ is orthogonal to the vector $(1,1,\ldots,1)$ when the neural state $\boldsymbol{x}(t)$ is nearly proportional to $(1,1,\ldots,1)$—which happens when all the options have equal values. $\mathcal{M}(t)$ is orthogonal to the vector $(1,1,\ldots,1,0)$ when $\boldsymbol{x}(t)$ is nearly proportional to $(1,1,\ldots,1,0)$ —which happens when we have $N-1$ equally high-valued options and a low-valued option

7. The uniqueness of the manifold over time: for two different time points $t$ and $t'$, the constraining manifolds $\mathcal{M}(t)$ and $\mathcal{M}(t')$ do not intersect with each other.

8. The diffusion process can be recovered by a renormalization: when we renormalize the neural activity $\boldsymbol{x}(t)$ by projecting it onto an $N-1$ dimensional hyperplane (the triangle in Fig. 2a) orthogonal to diagonal vector $(1,1,\ldots,1)$, the temporal evolution of population state on this plane is a standard $N-1$ dimensional diffusion process. Namely, the variance of temporal derivative of neural population activity is uniform over time in those dimensions.

9. The effect of opportunity cost: the position and speed of the constraining manifold $\mathcal{M}(t)$ depend not only on the number of options but also on the reward rate. This means that the offset activity of neurons should depend on the average reward size over trials or inter-trial interval, not only on the number of options.

##### Predictions about the neural states at the termination of evidence accumulation

Let $\boldsymbol{x}(t|\text{choose } i)$ denote the neural population state at the end of evidence accumulation, right before choosing option $i$.

10. The low-dimensional structure of the neural activity at the end of evidence accumulation: the neural activity when the stopping boundary is hit, $\boldsymbol{x}(t|\text{choose } i)$, is constrained on the $N-2$ dimensional manifold $B_i(t)$ (the 'end-point manifold') that is defined as the intersection of the constraining manifold $\mathcal{M}(t)$ and a hyper-plane, $x_i(t|\text{choose } i) = \theta_i$, where $x_i(t|\text{choose } i)$ is the $i$th compornent of $\boldsymbol{x}(t|\text{choose } i)$, and $\theta_i$ is a constant which is invariant to option sets.

11. The symmetry in the end-point manifolds: for different options $i$ and $j$, two end-point manifolds $B_i(t)$ and $B_j(t)$ do not intersect with each other. Moreover, on each $B_i(t)$, $x_j(t|\text{choose } i) < \theta_j$ for all $j \neq i$, and the distance between $B_1(t)$ and $B_2(t)$ is almost constant when the state $\boldsymbol{x}(t)$ is proportional to $(1,1,0,\ldots,0,0)$.

#### *Behavioral predictions*

12. The choice accuracy depends on time, the option set size, and the reward: the choice accuracy (the frequency of choosing the best options) decreases with reaction time. The choice accuracy also depends on the number of options as well as the reward rate.

13. The transitions of behavior between the ‘max-vs.-next’ and the ‘max-vs.-average’ strategy within a same task: in trials with $N$ almost equally-valued options, or with one high-valued option and $N-1$ low-valued options, the subject behavior (e.g., the reaction time dependency on choice contexts) is similar to what is predicted by the ‘max-vs.-average’ strategy (i.e., the strategy such that the decision is driven by the difference between the best option and the average of all the options.). In trials with two high-valued options and $N-2$ low-valued options, the subject behavior is similar to what is predicted by the ‘max-vs.-next’ strategy (i.e., the strategy such that the decision is driven by the difference between the best and the second-best options.). In other trials, the results differ from either of ‘max-vs.-average’ and ‘max-vs.-next’ strategies.

#### 4 Evidence with short- and long-range temporal correlations

Here we consider the modifications required to the optimal policy if the evidence features temporal short- and long-range correlations. In our discussion, we focus on positive temporal correlations. Similar arguments can be made for negative correlations. Both short- and long-range correlations reduce the amount of information available to the decision maker per unit time, but in different ways.

##### *Short-range correlations*

In the main text we have assumed the momentary evidence to be drawn i.i.d. according to $\delta \boldsymbol{x}_n | \boldsymbol{z} \sim \mathcal{N}(\boldsymbol{z}\delta t, \boldsymbol{\Sigma}_x \delta t)$, resulting in $\text{cov}(\delta \boldsymbol{x}_n, \delta \boldsymbol{x}_m) = \delta_{mn} \boldsymbol{\Sigma}_x \delta t$, where $\delta_{mn} = 1$ if $m = n$, and $\delta_{mn} = 0$ otherwise. That is, the momentary evidence provides independent information about $\boldsymbol{z}$ within each small time bin. This makes summing up this momentary evidence the optimal thing to do.

Let us now consider what happens if the momentary evidence becomes correlated across time. We first focus on short-range correlations, which could arise if the momentary evidence with white noise is passed through a circuit that low-pass filters this evidence. Then, we have $\text{cov}(\delta x_n, \delta x_{n+m}) > 0$ for sufficiently small time-differences $m\delta t$, and an autocorrelation that drops to zero with increasing $m\delta t$. What is the impact of these correlations on evidence accumulation and the optimal decision policy?

One effect of such correlations is that the amount of independent information about $\boldsymbol{z}$ per unit time is reduced. This is because consecutive pieces of momentary evidence are correlated, such that their associated noise does not average out when summing them. However, as long as the momentary evidence's auto-correlation structure is known and sufficiently well-behaved, we can apply a linear filter to the incoming momentary evidence to whiten it (e.g., Papoulis & Pillai, 2002[1]). This will result in another stream of momentary evidence that has the overall same amount of information about $\boldsymbol{z}$, but whose individual pieces of evidence are independent across time. Thus, the re-formatted momentary evidence satisfies the assumptions underlying the model developed in the main text, such that its conclusions still apply.

What are the limitations of this approach? First, temporal whitening of the momentary evidence requires knowledge of its auto-correlation structure. Knowing the statistics of the incoming evidence in a general pre-requisite to finding the optimal stopping boundaries, as finding them involves computing an expectation over potential future values of the accumulated evidence. Without knowing these statistics, we would not be able to find optimal stopping boundaries. This also applies to the auto-correlation structure. Second, despite temporal correlations, the amount of information per unit time needs to remain constant. For the original i.i.d. momentary evidence, this was satisfied by a likelihood $p(\delta \boldsymbol{x}_n | \boldsymbol{z})$ that had a fixed covariance structure. For correlated momentary evidence, this remains satisfied as long as its auto-correlation structure does not vary across time. Similar approaches to identifying optimal stopping boundaries can also be applied to scenarios in which the informativeness of momentary evidence fluctuates across time, but the resulting policies will become significantly more complex (e.g., Drugowitsch et al. , 2014[2]).

##### *Long-range correlations*

In the context of long-range correlations, we consider the case where the momentary evidence associated with each option is offset by a random, unknown, amount that is fixed within each trial. Let us denote this amount $y_j$ for option $j$ and assume it to be drawn independently in each trial from a zero-mean Gaussian with variance $\sigma_y^2$, that is $y_j \sim \mathcal{N}(0, \sigma_y^2)$. The momentary evidence in this trial is then drawn according to $\delta x_{j,n} | y_j, z_j \sim N(z_j + y_j, \sigma_x^2)$.

One extreme of this scenario are ballistic accumulator models[3,4], in which the only stochastic element is $y_j$, whereas the momentary evidence is noise-free, that is $\sigma_x^2 = 0$. In this case it becomes superfluous to accumulate evidence, as evidence samples beyond the first do not yield any additional information. Thus, the optimal policy would be to await this first sample and decide immediately after that.

For noisy momentary evidence, when $\sigma_x^2 > 0$, it remains optimal to accumulate momentary evidence. In this case, the only impact of an unknown $y_j$ is that the prior over the mean of the momentary evidence becomes less certain. Specifically, it grows in variance by $\sigma_y^2$. As a consequence, we can handle this case by ignoring the $y_j$'s, while at the time widening the prior over $z$ by $\sigma_z^2$. Therefore, it reduces to the case discussed in the main text, and results in the same optimal policy.

In summary, both short- and long-term temporal correlations in the momentary evidence reduce the amount of evidence we have about the true values of the underlying latent states $\boldsymbol{z}$. Short-term correlations do so by reducing the information in each piece of evidence, effectively making the likelihood less certain. Long-term correlations, in contrast, make the underlying mean less certain, effectively making the prior less certain. Both cases thus impact particular parameters of the model, while leaving the general structure unchanged. Therefore, while they impact the optimal decision boundaries quantitatively, they don't change our conclusions qualitatively.

#### 5 Discrete implementation of circuit with divisive normalization

In this section, we derive equations for our circuit model that approximates the optimal policy. Our network operates at two time-scales. On the slower time-scale, neurons accumulate (noisy) momentary evidence independently across options according to:

$$\boldsymbol{x_t} = C_t \,\delta \boldsymbol{x}_t + \frac{C_t}{C_{t-1}} \boldsymbol{x}_{t-1} \tag{30}$$

where $\boldsymbol{x_t}$ is the vector of accumulated evidence at time $t$, $\delta \boldsymbol{x}_t \sim \mathcal{N}(\boldsymbol{z}_t dt, \Sigma_t dt)$ is the vector of momentary evidence at time $t$, where $\boldsymbol{z}_t$ is the vector of "true" rewards, $\Sigma_t$ is a covariance matrix that makes the momentary evidence noisy, and $C_t$ is the commonly used divisive normalization term[5]:

$$C_t = \frac{K}{\sigma_h + \sum_{n=1}^{N} x_n(t)} \tag{31}$$

On the faster time scale, activity is projected onto a manifold defined by $\frac{1}{N}\sum_i f(x_i) = u(t)$, (shown as a gray surface in **Supplementary Figure S2**) where $u(t)$ is the urgency signals. This operation is implemented by iterating:

$$x_i \leftarrow x_i + \gamma \left( u(t) - \frac{1}{N} \sum_i f(x_i) \right) \tag{32}$$

until convergence, where $\gamma$ is the update rate and $f$ is a rectified polynomial non-linearity. Ignoring the fast dynamics, Equation (30) can be rearranged to get:

$$\boldsymbol{x_t} - \boldsymbol{x}_{t-1} = C_t \,\delta \boldsymbol{x}_t + \frac{C_t - C_{t-1}}{C_{t-1}} \boldsymbol{x}_{t-1} \tag{33}$$

$$d\boldsymbol{x}_t = C_t \left( \boldsymbol{z}_t dt + \Sigma_x^{\frac{1}{2}} dW_t \right) + \frac{dC_t}{C_t} \boldsymbol{x}_t \tag{34}$$

$$\frac{dx_j(t)}{dt} = C(t) \delta \boldsymbol{x}_t + \frac{1}{C(t)} \frac{dC(t)}{dt} x_j(t) \tag{35}$$

From Equation (31), it can be shown that

$$\frac{dC(t)}{dt} = -\frac{C^2(t)}{K} \sum_{n=1}^{N} \frac{dx_n(t)}{dt} \tag{36}$$

Combining Equations (35) and (36), we get

$$\frac{dx_j(t)}{dt} = C(t) \delta \boldsymbol{x}(t) - \frac{C(t)}{K} x_j(t) \sum_{n=1}^{N} \frac{dx_n(t)}{dt} \tag{37}$$

Equation (37) captures the slow dynamics of our circuit model. On a faster time-scale $\tau \ll dt$, this activity is further projected on to the manifold $\frac{1}{N}\sum_i f\big(x_i(t)\big) = u(t)$, which can be expressed as:

$$\frac{dx_j(t)}{dt} = C(t)\left(\delta\boldsymbol{x}(t) + \frac{1}{\tau}\left(u(t) - \frac{1}{N}\sum_{n=1}^{N} f\left(\frac{x_n(t)}{C(t)}\right)\right) - \frac{1}{K}\, x_j(t)\sum_{n=1}^{N}\frac{dx_n(t)}{dt}\right) \tag{38}$$

This can be written in vector form as:

$$\frac{1}{C_x}\frac{d\boldsymbol{x}(t)}{dt} = \delta\boldsymbol{x}(t) + \frac{1}{\tau}\left(u(t) - \frac{1}{N}\Big(\mathbb{1}\cdot f\big(\boldsymbol{x}(t)\big)\Big)\right) - \boldsymbol{x}(t)\left(\mathbb{1}\cdot\frac{d\boldsymbol{x}(t)}{dt}\right) \tag{39}$$

$$= \delta\boldsymbol{x}(t) + \frac{1}{\tau}\left(u(t) - \frac{1}{N}\Big(\mathbb{1}\cdot f\big(\boldsymbol{x}(t)\big)\Big)\right) - \boldsymbol{x}(t)\left(\frac{C_x\big(\mathbb{1}\cdot\tilde{\mathbf{z}}(\mathrm{t})\big)}{K + C_x\big(\mathbb{1}\cdot\boldsymbol{x}(t)\big)}\right) \tag{40}$$

where the explicit Equation (40) follows by summing Equation (39) and substituting the last term on its right-hand side.

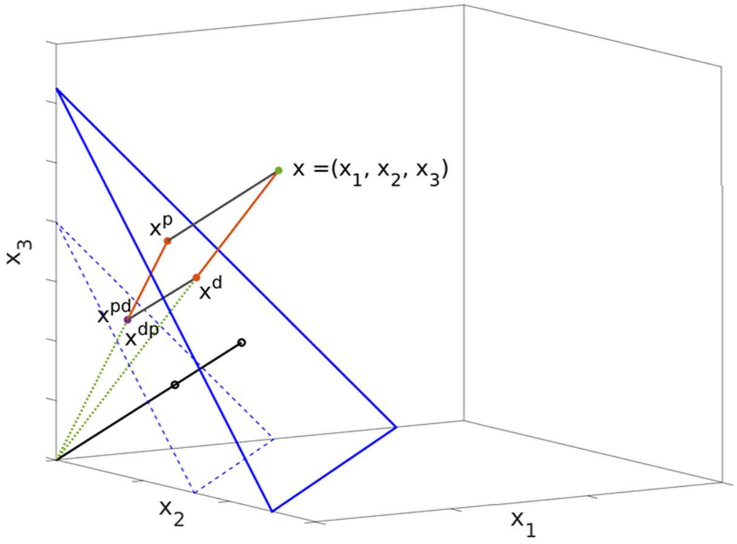


**Supplementary Figure S3**

Geometric depiction of the projection due to the linear constraint $(u(t) - \frac{1}{N}\sum_n^N x_n(t) = 0)$ and divisive normalization. The nonlinear constraint $(u(t) - \frac{1}{N}\sum_n^N f(x_n(t)) = 0)$ makes the blue triangular plane a curved surface. $\boldsymbol{x}$ is an arbitrary initial point. $\boldsymbol{x}^p$ is obtained by projecting $\boldsymbol{x}$ according to the nonlinear constraint mentioned above. Further applying divisive normalization to $\boldsymbol{x}^p$ gives $\boldsymbol{x}^{pd}$, *i.e.* $\boldsymbol{x}^{pd} = C\boldsymbol{x}^p$, where $C$ is defined in Eq. 31. On the other hand, $\boldsymbol{x}^d$ is obtained by applying divisive normalization to $\boldsymbol{x}$, *i.e.* $\boldsymbol{x}^d = C\boldsymbol{x}$, and further projecting $\boldsymbol{x}^d$ according to the nonlinear constraint mentioned above gives $\boldsymbol{x}^{dp}$. In the following section, we analytically show that $\boldsymbol{x}^{dp} = \boldsymbol{x}^{pd}$, or that the diffusion is unaffected if one were to apply the nonlinear constraint first followed by divisive normalization or *vice versa*.

##### *Optimal evidence accumulation*

Our full model which approximates the optimal policy by accumulating evidence and then projecting it on a manifold at each time-step. If we rescale the entire evidence accumulation space after this process at each time-step, then the relative distances between the accumulators and decision threshold are preserved, leaving the choices optimal. Mathematically, divisive normalization rescales the evidence accumulation space. As we just argued, doing so after the projection preserves optimality.

However, different instances of the discrete implementation may reverse this order – one may perform the rescaling before projecting at each time-step. Fortunately, it is possible to show analytically that, at least in the linear case, i.e. when $f(x) = ax + b$, the order is irrelevant. For simplicity, but without loss of generality, we will show this for $f(x) = x$.

To formally analyze this problem, we make use of some geometric intuition (see **Supplementary Figure S3)**. In 3-dimensional evidence accumulation space, consider an arbitrary initial point, $\boldsymbol{x} = (x_1, x_2, x_3)$, that has not hit any decision boundary. If we were to apply divisive normalization to this point, we would get $\boldsymbol{x}^d = (x_1^d, x_2^d, x_3^d)$, and further applying the projection would yield $\boldsymbol{x}^{dp} = (x_1^{dp}, x_2^{dp}, x_3^{dp})$, where the subscripts $d$ and $p$ are defined respectively by the form of divisive normalization and projection on a linear or a non-linear manifold (along the diagonal) as noted later. On the other hand, if we were to project the initial point $\boldsymbol{x}$ first, that would give us $\boldsymbol{x}^p = (x_1^p, x_2^p, x_3^p)$, and then implementing divisive normalization would yield $\boldsymbol{x}^{pd} = (x_1^{pd}, x_2^{pd}, x_3^{pd})$. Our goal is to show that $\boldsymbol{x}^{dp} = \boldsymbol{x}^{pd}$.

In the linear case,

$$x_i^p = x_i + u - \frac{1}{N}\sum_{n=1}^{N} x_n \tag{36}$$

$$x_i^d = C_x x_i$$

$$= \frac{K\, x_i}{\sigma_h + \sum_{n=1}^{N} x_n} \tag{37}$$

Using these, we can calculate $x_i^{pd}$ as

$$x_i^{pd} = \frac{K\, x_i^p}{\sigma_h + \sum_{n=1}^{N} x_n^p}$$

$$= \frac{K\, x_i + Ku - \frac{K}{N}\sum_{n=1}^{N} x_n}{\sigma_h + \sum_{n=1}^{N} x_n + Nu - \sum_{n=1}^{N} x_n}$$

$$= \frac{K\, x_i + Ku - \frac{K}{N}\sum_{n=1}^{N} x_n}{\sigma_h + Nu}$$

and $x_i^{dp}$ as

$$x_i^{dp} = x_i^d + u^d - \frac{1}{N}\sum_{n=1}^{N} x_n^d$$

$$= \frac{K\, x_i}{\sigma_h + \sum_{n=1}^{N} x_n} + \frac{K\, u}{\sigma_h + \sum_{n=1}^{N} x_n} - \frac{\frac{K}{N}\sum_{n=1}^{N} x_n}{\sigma_h + \sum_{n=1}^{N} x_n}$$

$$= \frac{K\, x_i + Ku - \frac{K}{N}\sum_{n=1}^{N} x_n}{\sigma_h + \sum_{n=1}^{N} x_n}$$

$$= x_i^{pd} \qquad (\because \sum_{n=1}^{N} x_n = Nu \text{ after projection})$$

Thus, the order does not matter in the linear case.

Adding the nonlinearity does not allow a closed form solution. The projection now takes place iteratively as $\boldsymbol{x}(t) \leftarrow \boldsymbol{x}(t) + \alpha[u(t) - \frac{1}{N}\Sigma_n x_n(t)]$ until convergence. It is important to note that though the projection is on a non-linear manifold, the direction of the projection is parallel to the diagonal; only the magnitude of the distance is determined iteratively.

#### 6 Dependencies of optimal stopping bounds on task parameters

We systematically explore how task parameters depend on the optimal stopping bounds. In **Supplementary Figure S5**, we show how the decision boundaries change as a function of time (a), inter-trial interval (b), noise variance (c), and with symmetric (d) and asymmetric (e) prior mean of reward. We have not been able to derive analytical approximations to the stopping bounds but note that the neural network provides a close approximation to the optimal bound with only three parameters. Given the shape and time dependence of the bounds, it is unlikely that it is possible to obtain an analytical solution with fewer parameters.

#### 7 Additional Supplementary Figures

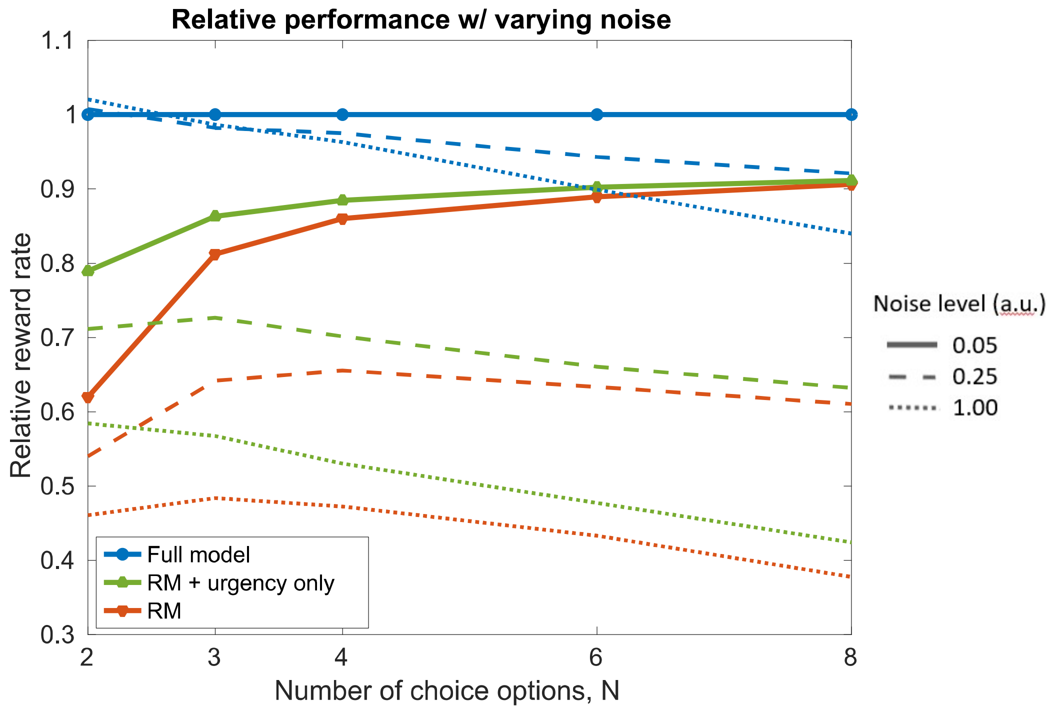


**Supplementary Figure S4. Decision-bound variability affects models' relative performance**

The race model variants without constrained evidence accumulation approximating the optimal policy perform much worse than our model's variants with that constraint, a result that is demonstrated in **Figure 5c**. Here, we show that reducing the amount of variability in the decision bounds brings the models' relative performances closer to each other as was the case in **Figure 2d**.

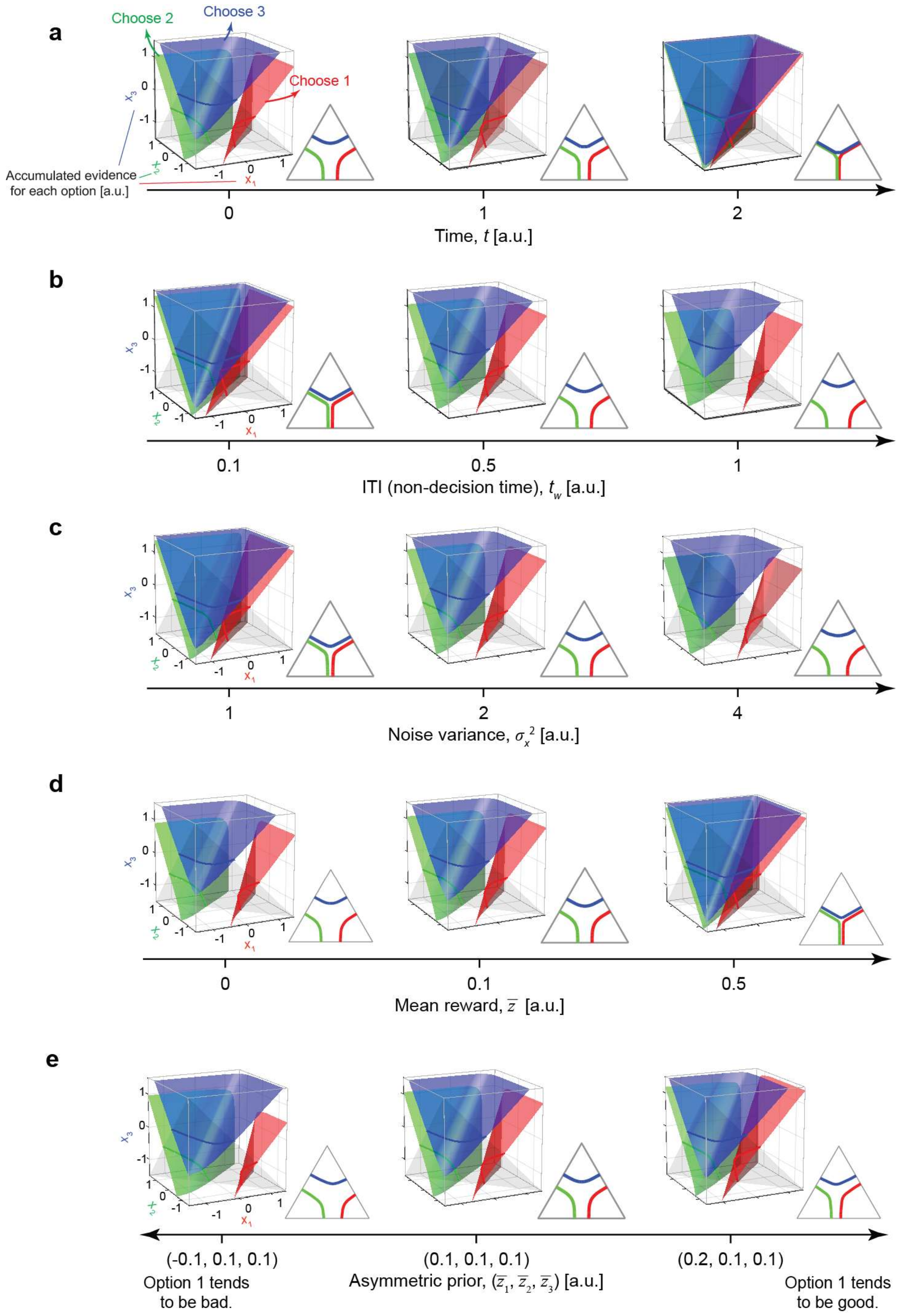


**Supplementary Figure S5. Dependencies of the stopping boundaries on task parameters.**

(**a**) Dynamics of decision boundaries over time, $t$. The decision boundaries approach each other over time. Here, we used the following parameters: reward prior, $(\bar{z}_1, \bar{z}_2, \bar{z}_3) = (\bar{z}, \bar{z}, \bar{z}) = (0.1, 0.1, 0.1)$; inter trial interval (ITI, including non-decision time), $t_w = 0.5$; noise variance, $\sigma_x^2 = 2$.

In (**b**)-(**e**) we varied a single parameter, while keeping all other parameters constant. The shown boundaries are the initial ones, at time $t = 0$.

(**b**) Effect of inter trial interval (ITI), $t_w$. The boundaries start further apart for longer ITIs . $t_w = 0.5$ corresponds to the leftmost plot in panel **a**.

(**c**) Effect of the evidence noise variance, $\sigma_x^2$. The boundaries start further apart for larger noise. $\sigma_x^2 = 2$ corresponds to the leftmost plot in panel **a**.

(**d**) Effect of the reward prior mean, $\bar{z}$. The boundaries start closer to each other for larger mean rewards. $\bar{z} = 0.1$ corresponds to the leftmost plot in panel **a**.

(**e**) Effect of the asymmetric reward prior, $(\bar{z}_1, \bar{z}_2, \bar{z}_3)$, where $\bar{z}_1$, $\bar{z}_2$, and $\bar{z}_3$ can be different from each other. The boundaries remain parallel to the cube diagonal but the asymmetric priors cause a shift of the boundary positions when projected on the triangle orthogonal to the diagonal, such that the boundaries corresponding to the most rewarded options start closer to the center of the triangle. $(\bar{z}_1, \bar{z}_2, \bar{z}_3) = (0.1, 0.1, 0.1)$ is identical to the leftmost plot in panel **a**.

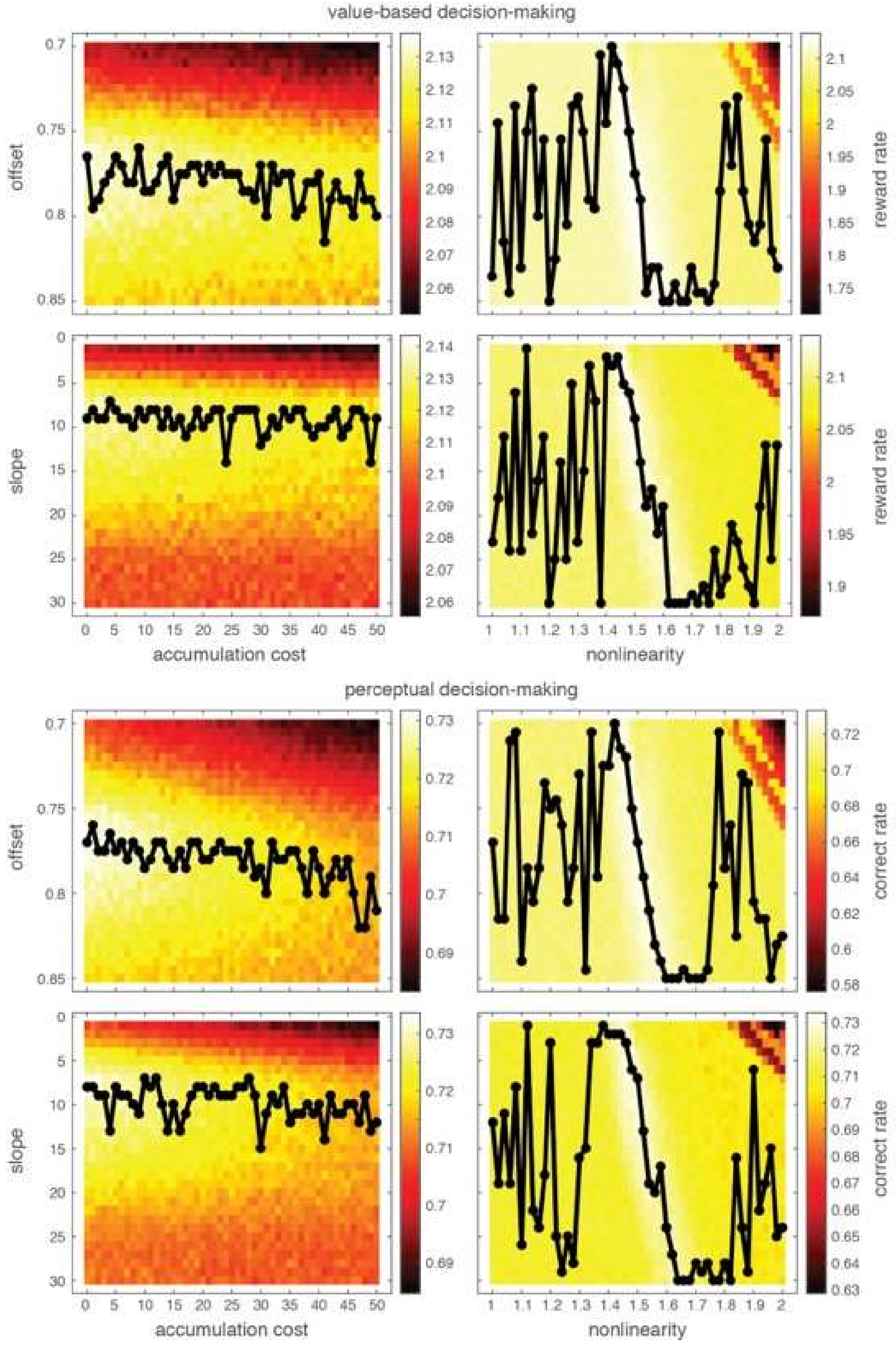
value-based decision-making
offset
slope
reward rate
accumulation cost
nonlinearity
perceptual decision-making
offset
slope
correct rate
accumulation cost
nonlinearity

**Supplementary Figure S6. The optimal urgency signal is only weakly dependent on accumulation cost and nonlinearity.** Each panel shows for combinations of urgency signal parameters (vertical axis; offset or slope) and cost (left panels) or nonlinearity setting (right panels) the reward rate (value-based decisions; top) or correct rate (perceptual decision; bottom) as a color gradient. For each parameter combination, reward and correct rate were found by simulating 500.000 trials. The black line in each panel indicates for each cost or nonlinearity setting the value of the urgency signal parameter that maximizes the reward/correct rate. This line is noisy due to the simulation-based stochastic evaluation of the reward/correct rates. In general, both optimal slope and offset only weekly depend on the accumulation cost. The same applies to the nonlinearity, except for a narrow band around 1.5, where it is best to decrease both slope and offset for an increase in this nonlinearity.

#### References


1. Papoulis, A. & Pillai, S. Probability, random variables, and stochastic processes. (2002).
2. Drugowitsch, J., DeAngelis, G., Klier, E., Elife, D. A.- & 2014, undefined. Optimal multisensory decision-making in a reaction-time task. *cdn.elifesciences.org*
3. Carpenter, R. & Williams, M. Neural computation of log likelihood in control of saccadic eye movement. *Nature* **377,** 59–62 (1995).
4. Brown, S. & Heathcote, A. A Ballistic Model of Choice Response Time. *Psychol. Rev.* **112,** 117–128 (2005).
5. Churchland, A. K. & Ditterich, J. New advances in understanding decisions among multiple alternatives. *Curr. Opin. Neurobiol.* **22,** 920–926 (2012).
